# The expanding avian influenza panzootic: skua die-off in Antarctica

**DOI:** 10.1101/2025.04.25.650384

**Authors:** Matteo Iervolino, Anne Günther, Lineke Begeman, Begoña Aguado, Theo M. Bestebroer, Beatriz Bellido-Martin, Adam Coerper, M. Valentina Fornillo, Bruno Fusaro, Andrés E. Ibañez, Lonneke Leijten, Simeon Lisovski, Mariané B. Mañez, Alice Reade, Peter van Run, Florencia Soto, Ben Wallis, Meagan Dewar, Antonio Alcamí, Martin Beer, Ralph E.T. Vanstreels, Thijs Kuiken

**Affiliations:** Department of Viroscience, Erasmus University Medical Center; Rotterdam, the Netherlands; Institute of Diagnostic Virology, Friedrich-Loeffler-Institut, Federal Research Institute for Animal Health; Greifswald-Insel Riems, Germany; Centro de Biología Molecular Severo Ochoa, Consejo Superior de Investigaciones Científicas (CSIC) and Universidad Autónoma de Madrid (UAM); Madrid, Spain; Ocean Expeditions Support Vessel S/V Australis; Sydney, New South Wales, Australia; Argentine Antarctic Institute, Center for Parasitological and Vector Studies; La Plata, Argentina; Sección Ornitología, Div. Zool. Vert. Museo de la Plata (FCNyM-UNLP-CONICET); La Plata, Argentina; Alfred Wegener Institute Helmholtz Centre for Polar and Marine Research; Potsdam, Germany; Instituto de Biología de Organismos Marinos (IBIOMAR-CONICET); Puerto Madryn, Argentina; Future Regions Research Centre, Federation University Australia; Ballarat, Victoria, Australia; Karen C. Drayer Wildlife Health Center, One Health Institute, School of Veterinary Medicine, University of California, Davis; Davis, CA, USA

## Abstract

High pathogenicity avian influenza virus of subtype H5 (H5 HPAIV), clade 2.3.4.4b, invaded Antarctica in 2023. Here we show that H5 HPAIV caused high mortality in a breeding colony of skuas at one of ten sites we visited in March 2024. By combined virological and pathological analyses, we found that H5 HPAIV caused multi-organ necrosis and rapid death in skuas. Taken together with recent data, skuas in Antarctica are at risk of continued mortality from H5 HPAIV infection, threatening their already small populations. Conversely, because of their wide distribution and ecological relevance, skuas may play a substantial role in spread of the virus across Antarctica. Transdisciplinary surveillance is needed in coming years to monitor the impact of this poultry-origin disease on Antarctica’s unique wildlife.

## Introduction

The incursion of A/goose/Guangdong/1/96 (Gs/Gd) lineage high pathogenicity avian influenza (HPAI) virus (HPAIV) of the subtype H5 clade 2.3.4.4b into Antarctica is a potential threat to the millions of wild birds and mammals living there. This is exemplified by the incursion of H5 HPAIV into South America in 2022-2023, where it is estimated to have killed at least 667,000 wild birds of 83 species and 53,000 wild mammals of 11 species (*1, 2*). There is also a real risk that the mortality of Antarctic wildlife from this virus will go unnoticed or underreported. Such documentation is critical to support the need for transformative change to protect wildlife populations from HPAI and other anthropogenic diseases originating from ever-increasing livestock populations *(3, 4)*.

The effects of H5 HPAIV infection on wildlife range from no clinical signs to death (*5*). This means that infected wild animals can both act as vectors of the virus, possibly carrying it thousands of kilometers during long-distance migration in a staggered manner (*6*), and as its victims, suffering severe die-offs (*7*). It also means that detection of HPAIV in a dead wild animal needs to be complemented by pathological demonstration of virus-associated lesions to confirm that HPAI virus infection was the cause of death. Additionally, these investigations are enhanced by testing apparently healthy individuals in wildlife populations.

The presence of H5 HPAIV of clade 2.3.4.4b in Antarctica was first recorded in January 2024 in a kelp gull (*Larus dominicanus*) on Livingston Island (South Shetland Islands), adjacent to the Antarctic Peninsula (*8, 9*). The virus may have spread to Antarctica directly from the southern tip of South America (*8*), or it may have used Subantarctic islands as stepping stones (*10*). Seabirds and marine mammals are known to migrate between these locations during their annual cycle and may have carried the virus with them (*11, 12*).

Two closely related seabird species that are known to be susceptible to H5 HPAIV infection in the southern hemisphere are brown skuas (*Catharacta antarctica*) and south polar skuas (*Catharacta maccormicki*) (*10, 13*). While brown skuas, with a global population of about 7000 pairs, exhibit a circumpolar breeding and non-breeding distribution across the southern hemisphere (*14-16*), south polar skuas, with a global population of 5000 to 7000 pairs, breed around the coast of Antarctica (*14, 17*), but exhibit a non-breeding distribution on the North Pacific and North Atlantic Oceans (*18, 19*). In some locations, these species breed in sympatry, and hybridization occurs (*15*). Both species were among the first to be found infected with H5 HPAIV in Subantarctic South Georgia in late 2023 as well as Antarctica in early 2024 (*10, 13, 20, 21*). Although H5 HPAIV was detected in association with morbidity and mortality in both brown and south polar skuas, no pathological analyses have been presented to confirm HPAI as the cause of death.

Besides HPAI, avian cholera needs to be considered in the differential diagnosis of unusual mortality of wild birds in the Subantarctic and Antarctic regions (*22-25*). *Pasteurella multocida*, the causative agent of avian cholera, typically causes epidemics in wetlands or breeding colony sites with high densities of birds, including seabirds (*26*).

Little is known about the spread of H5 HPAIV within and among species of wild seabirds and mammals in Antarctica, about the character and severity of disease, and the levels of mortality in affected populations. Therefore, we set up the HPAI Australis Expedition to investigate the introduction and spread of H5 HPAIV across the Antarctic Peninsula and its impact on local wildlife (*20*). Between 17 and 28 March 2024, at the end of the austral summer, when the breeding season of skuas, gentoo penguins (*Pygoscelis papua*) and Adélie penguins (*Pygoscelis adeliae*) had already concluded, we conducted epidemiological surveys at 10 locations at the South Shetland Islands, Weddell Sea and Trinity Peninsula for wildlife morbidity and mortality, performed autopsies and collected postmortem tissue samples and environmental samples for virological, bacteriological, and pathological analyses. Here, we report our findings of H5 HPAIV detection in skuas at three locations (Hope/Esperanza Bay, Devil Island, Beak Island) at the Trinity Peninsula and Weddell Sea region, and confirmation of HPAI as the probable cause of a mass die-off of south polar skuas at one of those locations (Beak Island). Additionally, we present results from avian cholera, environmental samples and other species from which samples were collected.

## Results

### Overview of skua mortality during the HPAI Australis Expedition

During our expedition (Fig. 1), we found evidence of skua mortality at six sites (Hope Bay, Devil Island, Beak Island, Heroina Island, Beagle Island and Paulet Island), four of which are presented here (results for Heroina and Beagle Island will be shown elsewhere) (Fig. 2, Table 1).

**Table 1.** Overview of diagnostic criteria for high pathogenicity avian influenza (HPAI), avian cholera, and fungal infection as proposed causes of death of 20 skuas found dead. The confidence level of the diagnosis applies to all diagnoses within the same animal.

| Location | Individual | Species | Age class | Nutritional condition | HPAI |  | Avian cholera |  | Fungal disease | Diagnosis |  |
| --- | --- | --- | --- | --- | --- | --- | --- | --- | --- | --- | --- |
|  |  |  |  |  | Brain H5 HPAIV RNA | Viral histo | Brain *P.m.* DNA | *P.m.* histo | Fungus histo | Diseases at death in order of severity | Confidence level |
| Hope Bay | SK01 | BRSK | Ad | M | - | - | +++ | + | + | 1. Fungal disease<br>2. Avian cholera | High |
|  | SK02 | BRSK | Ad | P | ++ | na | ++ | na | na | 1. Avian cholera<br>2. HPAI | Medium |
|  | SK03 | SPSK | Ad | VP | + | na | +++ | na | na | 1. Avian cholera<br>2. HPAI<br>3. Starvation | Medium |
|  | SK04 | UNSK | Ad | VP | - | na | ++ | na | na | 1. Avian cholera<br>2. Starvation | Medium |
|  | SK05 | UNSK | Ad | P | - | na | ++ | na | na | 1. Avian cholera | Medium |
|  | SK06 | UNSK | Ad | M | - | - | +++ | + | - | 1. Avian cholera | High |
| Devil Is. | SK07 | UNSK | Juv | G | +++ | + | - | - | - | 1. HPAI | High |
| Beak Is. | SK08 | SPSK | Ad | VP | +++ | + | - | - | + | 1. HPAI<br>2. Fungal disease<br>3. Starvation | High |
|  | SK09 | UNSK | Juv | VP | +++ | + | - | - | - | 1. HPAI<br>2. Starvation | High |
|  | SK10 | UNSK | Juv | G | +++ | + | - | - | - | 1. HPAI | High |
|  | SK11 | UNSK | Juv | G | +++ | + | - | - | - | 1. HPAI | High |
|  | SK12 | UNSK | Juv | G | +++ | + | - | - | - | 1. HPAI | High |
|  | SK13 | UNSK | nd | nd | +++ | na | - | na | na | 1. HPAI | Medium |
|  | SK14 | UNSK | nd | nd | +++ | na | - | na | na | 1. HPAI | Medium |
|  | SK15 | UNSK | nd | nd | ++ | na | - | na | na | 1. HPAI | Medium |
|  | SK16 | UNSK | nd | nd | +++ | na | - | na | na | 1. HPAI | Medium |
|  | SK17 | UNSK | nd | nd | +++ | na | - | na | na | 1. HPAI | Medium |
| Paulet Is. | SK29 | UNSK | Ad | nd | - | - | +++ | + | - | 1. Avian cholera | High |
|  | SK30 | UNSK | Ad | P | (+) | - | ++ | + | + | 1. Fungal disease<br>2. Avian cholera | High |
|  | SK31 | UNSK | Ad | na | - | - | +++ | na† | - | 1. Avian cholera | Medium |

**Figure 1.**
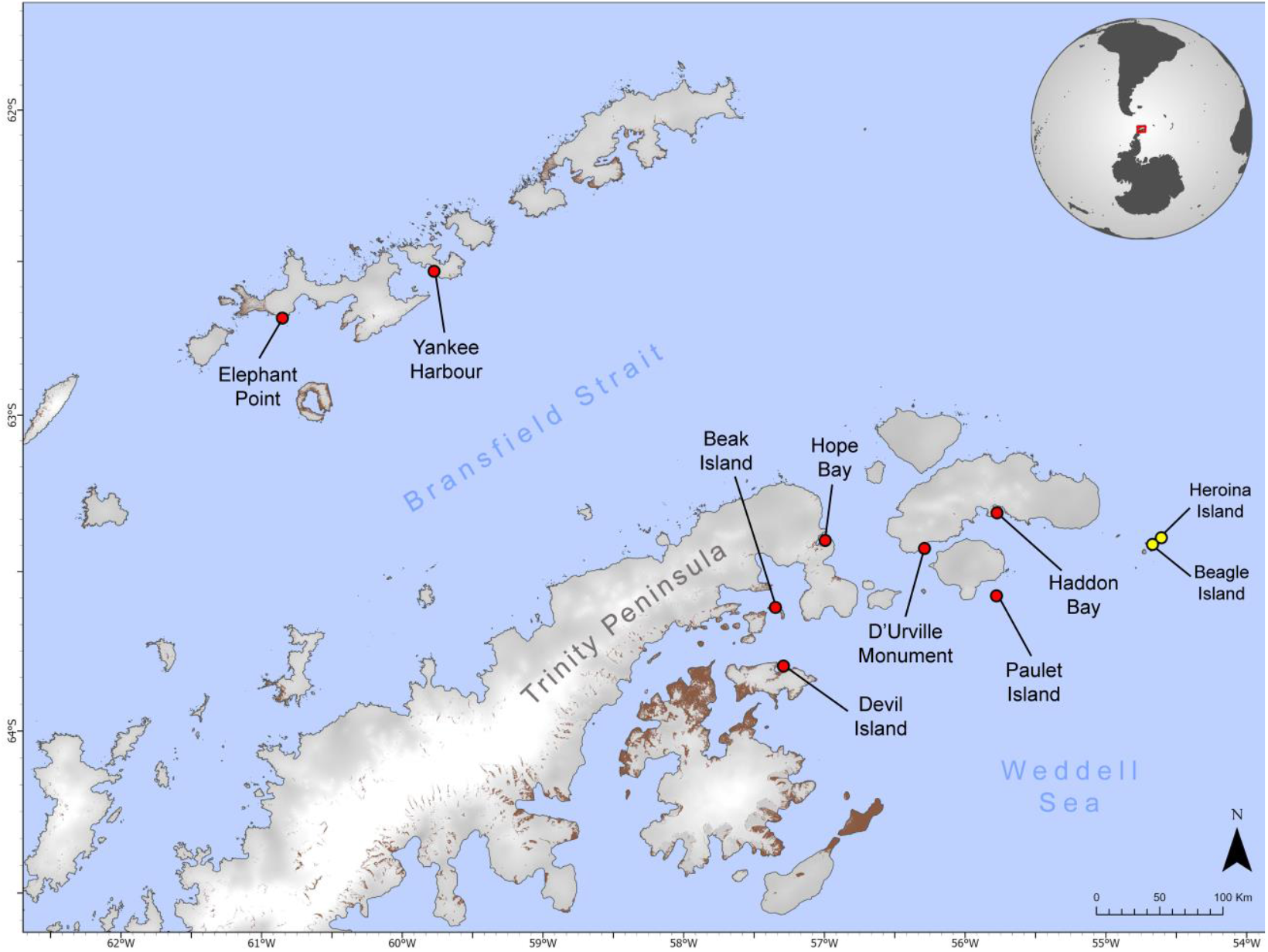
Map of the HPAI Australis Expedition, indicating investigated sites. Results from Heroina Island and Beagle Island (yellow) will be shown elsewhere.

**Figure 2.**
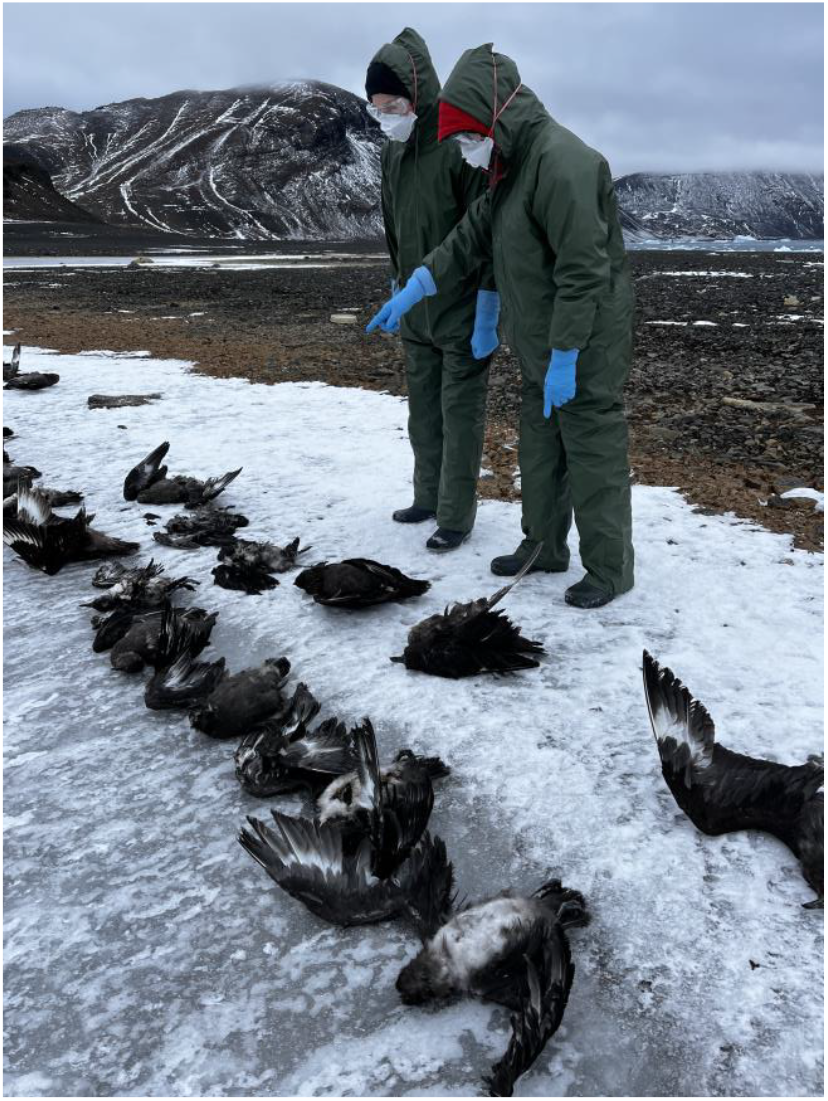
Dead skuas on Beak Island being examined by two expedition members. The individuals depicted in the photograph are co-authors of this manuscript and have consented to the inclusion of their images.

At Hope Bay, 16 non-breeding brown skuas and 3 non-breeding south polar skuas were previously found dead between February 5th and March 9th, 2024, through routine surveillance conducted by the Argentine National Antarctic Program. At the time of our visit, we could retrieve 6 of these dead adult skuas (1 south polar, 2 brown, 3 unidentified). Additionally, about 60 live skuas (apparently healthy) were present at the time of our visit. We necropsied two skuas and diagnosed avian cholera as the probable cause of death (SK01, SK06), although in one (SK01) a fungal infection probably contributed to death. We detected moderate to high levels of *P. multocida* in the brains of four additional dead skuas (SK02 to SK05). Two of these also had low (SK03) to moderate (SK02) levels of H5 HPAIV RNA in the brains (Table 2), so the cause of death remained unsure.

**Table 2.** H5 HPAIV viral loads (40-Cq, *M1* gene) and *P. multocida* (P.m.) bacterial loads (40-Cq, *kmt1* gene) in swabs and tissues of skuas found dead. Spleen, kidney and feather follicle were not tested for the presence of *P. multocida* DNA. The IDs from onboard testing are indicated to compare with previous results *(20, 40)*.

| Location | Individual | ID onboard test (40) | Tracheal swab |  | Oropharyngeal swab |  | Cloacal swab |  | Brain |  | Lung |  | Intestine |  | Liver |  | Spleen | Kidney | Feather follicle |
| --- | --- | --- | --- | --- | --- | --- | --- | --- | --- | --- | --- | --- | --- | --- | --- | --- | --- | --- | --- |
|  |  |  | HPAIV | *P. m.* | HPAIV | *P. m.* | HPAIV | *P. m.* | HPAIV | *P. m.* | HPAIV | *P. m.* | HPAIV | *P. m.* | HPAIV | *P. m.* | HPAIV | HPAIV | HPAIV |
| Hope Bay | SK01 | Skua_1_Hope | na | na | nd | 17.2 | nd | 16.8 | nd | 15.8 | nd | 18.5 | nd | 18.9 | nd | 4.1 | nd | nd | nd |
|  | SK02 | Skua_2_Hope | na | na | 10.6 | 4.1 | 7.4 | 9.9 | 18.6 | 9.7 | na | na | na | na | na | na | na | na | na |
|  | SK03 | Skua_3_Hope | na | na | 5.2 | 13.2 | 0.2 | 15.4 | 5.6 | 16.0 | na | na | na | na | na | na | na | na | na |
|  | SK04 | Skua_4_Hope | na | na | nd | 8.5 | nd | 4.9 | nd | 8.2 | na | na | na | na | na | na | na | na | na |
|  | SK05 | Skua_5_Hope | na | na | nd | 14.0 | nd | 14.2 | nd | 12.4 | na | na | na | na | na | na | na | na | na |
|  | SK06 | Skua_6_Hope | na | na | nd | 14.5 | nd | 16.1 | nd | 18.0 | nd | 19.9 | nd | 18.7 | nd | 22.3 | nd | nd | nd |
| Devil Is. | SK07 | Skua_1_Devil | na | na | 17.7 | nd | 17.9 | nd | 27.2 | nd | 26.7 | nd | 19.7 | nd | 21.0 | nd | 21.8 | 21.3 | 13.5 |
| Beak Is. | SK08 | Skua_1_Beak | nd | nd | na | na | 6.0 | nd | 20.5 | nd | 13.4 | nd | 7.9 | nd | 8.4 | nd | na | 6.6 | 2.2 |
|  | SK09 | Skua_2_Beak | 15.9 | nd | na | na | 12.2 | nd | 22.2 | nd | 18.4 | nd | 11.2 | nd | 14.6 | nd | 14.3 | 9.8 | 15.0 |
|  | SK10 | Skua_3_Beak | 16.8 | nd | na | na | 13.5 | nd | 21.4 | nd | 19.1 | nd | 15.5 | nd | 12.7 | nd | 13.7 | 14.8 | 9.5 |
|  | SK11 | Skua_4_Beak | 12.9 | nd | na | na | 9.4 | nd | 24.0 | nd | 18.0 | nd | 13.9 | nd | 10.4 | nd | 12.2 | 11.6 | 10.0 |
|  | SK12 | Skua_5_Beak | 19.8 | nd | na | na | 18.5 | nd | 25.0 | nd | 26.9 | nd | 23.8 | nd | 27.5 | nd | 27.0 | 25.3 | 15.6 |
|  | SK13 | Skua_6_Beak | na | na | na | na | na | nd | 28.0 | nd | na | nd | na | nd | na | nd | na | na | na |
|  | SK14 | Skua_7_Beak | na | na | na | na | na | nd | 21.7 | nd | na | nd | na | nd | na | nd | na | na | na |
|  | SK15 | Skua_8_Beak | na | na | na | na | na | nd | 16.3 | nd | na | nd | na | nd | na | nd | na | na | na |
|  | SK16 | Skua_9_Beak | na | na | na | na | na | nd | 22.2 | nd | na | nd | na | nd | na | nd | na | na | na |
|  | SK17 | Skua_10_Beak | na | na | na | na | na | nd | 25.0 | nd | na | nd | na | nd | na | nd | na | na | na |
| Paulet Is. | SK29 | Skua_1_Paulet | na | na | na | na | nd | na | nd | 18.5 | nd | 21.2 | na | na | na | na | na | na | na |
|  | SK30 | Skua_2_Paulet | na | na | na | na | nd | na | nd | 8.9 | nd | 18.6 | na | na | na | na | na | na | na |
|  | SK31 | Skua_3_Paulet | na | na | na | na | nd | na | nd | 15.1 | nd | 15.7 | na | na | na | na | na | na | na |

nd: not detected
na: not applicable (not sampled)

At Devil Island, we found a dead juvenile skua (unidentified species; SK07), and diagnosed HPAI as its probable cause of death. An additional 15 apparently healthy live skuas were present at the time of our visit.

We visited Beak Island because we considered its topography to provide a likely breeding habitat for skuas. There, we found 46 dead skuas (18 south polar, 28 unidentified species; 26 adult, 19 juvenile, 1 undetermined age class). Additionally, about 100 apparently healthy live skuas were also present at the time of our visit; these were occasionally seen scavenging on skua carcasses, in addition to agonistic interactions with other live skuas and kelp gulls. We necropsied five of the dead skuas (SK08 to SK12) and diagnosed HPAI as the most probable cause of death. We detected moderate (SK15) or high levels (SK13, SK14, SK16, SK17) of H5 HPAIV in the brains of five additional dead skuas, supporting HPAI as cause of death.

At Paulet Island, we found three dead skuas (unidentified species; SK29 to SK31) and we determined that the possible causes of death were avian cholera in two of them (SK29, SK31), and either fungal disease or avian cholera in the third (SK30). About 30 apparently healthy live skuas were present at the time of our visit.

### Macroscopic findings in skuas found dead

Some of the skuas that were found dead had abnormal postures, possibly related to HPAI: opisthotonos (backward arching of head, neck, and spine; 15/37), torticollis (twisting of the neck; 8/37) and/or wings spread (32/37; table S1, fig. S1). Out of 6 dead skuas (1 from Devil Island, 5 from Beak Island) diagnosed with HPAI as a probable or possible cause of death, one had opisthotonos (SK07), one had torticollis (SK08), and four had wings spread (SK08 to SK11). However, out of two dead skuas from Paulet Island (SK29 and SK30) diagnosed with avian cholera as a probable or possible cause of death, SK30 had opisthotonos and both had wings spread, indicating that these abnormal postures are not restricted to death from HPAI. The postures of the dead skuas from Hope Bay were not recorded as the carcasses had been previously manipulated to avoid scavenging by healthy skuas.

None of the 11 skuas examined had significant macroscopic lesions, with three exceptions. One skua diagnosed with HPAI as probable cause of death (SK07) had diffusely dark red and wet lungs (pulmonary oedema) (fig. S1C). Two skuas diagnosed with fungal disease as probable or possible cause of death (SK01, SK08) had multiple characteristic white, dull, firm, well demarcated foci (3 to 30 mm in diameter) on tracheal mucosa, lungs and heart. The nutritional conditions of the autopsied skuas ranged from very poor to good, and were not correlated with the diagnoses of HPAI, avian cholera, or fungal disease as causes of death (Table 1). All the autopsied skuas had empty stomachs, suggesting they had not fed recently before death or had regurgitated their last meal.

Nine dead skuas were tested on site for the presence of influenza A virus by use of a rapid antigen test on swabs from different tissues (table S1). Out of 5 skuas that were subsequently H5 HPAIV-negative by RT-qPCR (Table 1; SK01, SK06, SK29 to SK31), 4 tested negative by rapid antigen test (SK01, SK06, SK29 and SK30) and for 1 the test result was uncertain (SK31). Out of 4 skuas that were subsequently H5 HPAIV-positive by RT-qPCR (SK02, SK07, SK09 and SK10; Table 1), 1 tested positive by rapid antigen test (SK07), 1 tested negative (SK02) and for 2 the test results were uncertain (SK09 and SK10).

### Virological and bacteriological analyses of swabs and tissues and serological analyses from skuas found dead

Influenza A virus RNA was detected by RT-qPCR from swabs and tissues of sampled skuas. H5 HPAIV was detected in swabs and tissues of 13 of 20 dead skuas tested, based on the combination of positive M1, H5, and HPAI RT-qPCRs for each tissue or swab (Table 2, table S2). RNA values (40 – Cq) of the GAPDH housekeeping gene ranged between 11.7 to 25.0 units (19.6 ± 3.3 on average), indicating an overall good RNA preservation in the carcass samples (table S9). To confirm the presence of the multi-basic cleavage site (MBCS), a region of the hemagglutinin gene was sequenced from at least one sample from each H5 HPAIV-positive skua (table S2). Samples from all positive skuas were confirmed to have the same MBCS (PLREKRRKR/GLF). For each of the 13 skuas that tested positive, all sampled tissues and swabs tested positive by the three RT-qPCRs - indicating systemic H5 HPAIV infection - except for the tracheal swab of one dead skua (SK08). The highest viral RNA loads were consistently measured in the brain, except for one skua (SK12), in which the liver had the highest viral RNA load. Average viral RNA loads (expressed as 40 – Cq) were several units higher in the brain (21.4 ± 5.7) than in tracheal swabs (16.3 ± 2.9) and oropharyngeal swabs (11.1 ± 6.3). The dead skuas sampled at Hope Bay had viral RNA loads in the brain (5.6 and 18.6) that were substantially lower than the dead skuas sampled at Devil Island and Beak Island (23.1 ± 3.3 on average). Overall, the highest viral loads in skuas that were diagnosed with HPAI as a possible cause of death (SK07 to SK12) were found in the brain (23.4 ± 2.5 on average) (Fig. 3).

**Figure 3.**
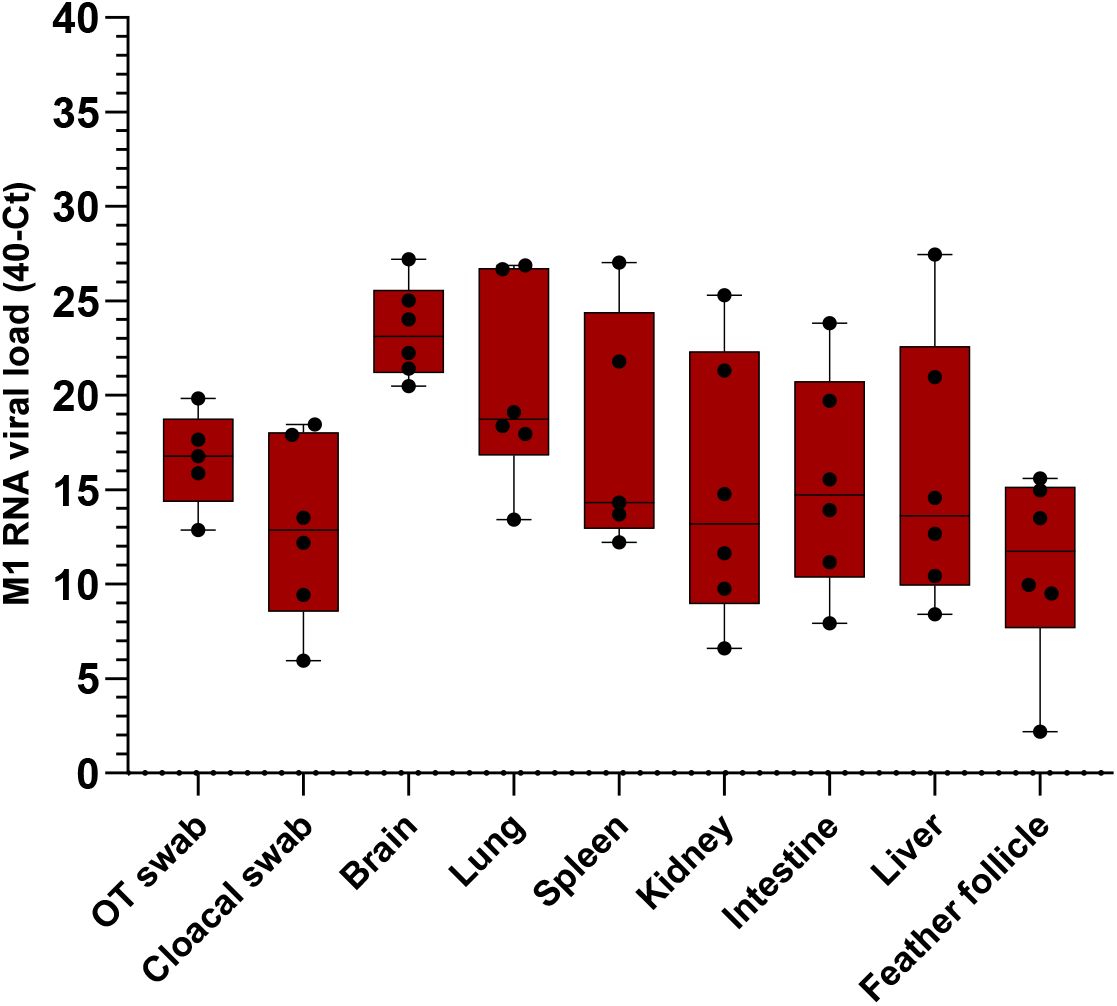
Influenza A RNA viral loads in different organs of skuas. Boxplot of the relative expression of M1 viral RNA loads (40 – Cq) in different organs and tissue swabs (OT swab: oropharyngeal or tracheal swab) of skuas diagnosed with HPAI as cause of death and tested positive for M1, H5 and HPAI qPCRs (SK07-12). Each black dot represents a single detection.

No antibodies against H5 influenza A viruses or nucleoprotein (NP) were detected by competitive ELISA in serum samples collected from two skuas (SK07 and SK12), both of which were diagnosed with HPAI as probable cause of death.

To determine whether or not avian cholera contributed to mortality, all the skuas were tested for the presence of *P. multocida* DNA (*kmt1* gene) by qPCR. *P. multocida* DNA was detected in swabs and tissues from all dead skuas in Hope Bay and Paulet Island (Table 2). The qPCR amplicon sequencing revealed up to 100% identity with *P. multocida* strains via BLASTn search (GenBank accession code pending). The *P. multocida* bacterial DNA load (40 – Cq) differed substantially between skuas, ranging from 8.2 to 18.5 units in the brain. In contrast, none of the tested organs or swabs from any of the skuas from Beak Island or Devil Island tested positive for *P. multocida* DNA by qPCR (Table 2).

### Histopathology of tissues from skuas found dead

#### Virus antigen expression in H5 HPAIV RNA-positive skua carcasses

Influenza A virus NP antigen was detected in all (n=6) H5 HPAIV RNA-positive skua carcasses (table S3). Brain was the organ that most consistently expressed virus antigen (5/6 skuas). Other tissues in which antigen expression was detected in parenchymal cells were lung, pancreas, thyroid, adrenal, preen gland and gonad (testicle) (fig. S2). One bird (SK12) had abundant antigen expression in endothelial cells in every tissue examined. Antigen was rarely detected in parenchymal cells of kidneys (1/6), and not in those of livers (0/6), or in hearts (0/6).

In the brain, virus antigen was detected in cerebrum, cerebellum and brainstem in a pattern that was seemingly random and differed per bird. Virus antigen also was detected in peripheral ganglia of one bird (fig. S2). The cells that expressed antigen in the nervous tissues were both neurons and glial cells, based on morphology and their location relative to other cells. In one bird (SK07), a large stretch of ependymal cells and the underlying neuropil also expressed virus antigen, suggesting entry via the cerebrospinal fluid (fig. S2). By *in situ* hybridization, we detected influenza A virus RNA in the brain section of a skua (SK11) that was virus antigen negative by immunohistochemistry (supplementary text S1, fig. S4).

In the lung (2/6), virus antigen was detected in air capillaries. Because of their close alignment, it was not possible to determine whether endothelial cells and/or epithelial cells of the air capillaries expressed virus antigen. Virus antigen distribution was multifocal to coalescing and involved 90 to 100% of the air capillaries per lung section.

In other tissues, the cell types expressing virus antigen were exocrine glandular epithelial cells in pancreas (Fig. 4) and epithelial cells of the preen gland, sperm producing cells (spermatogonia) of inactive testis, epithelial cells in thyroid gland (fig. S2) and in adrenal gland. In most tissues, virus antigen was mostly expressed by clusters of tens of cells.

**Figure 4.**
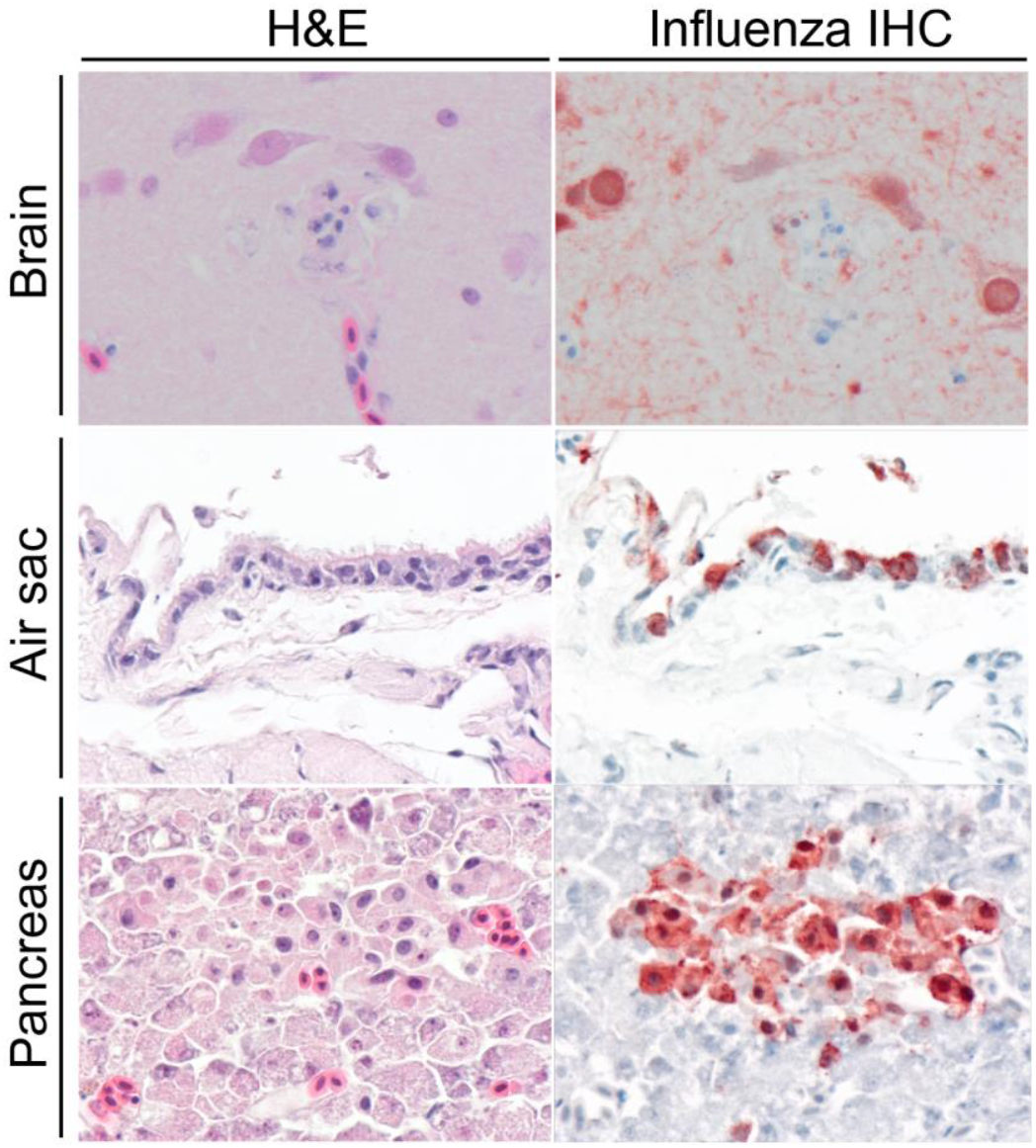
Serial sections of hematoxylin and eosin staining (H&E) and influenza A NP IHC in tissues of skuas found dead. Brain and pancreas belong to SK07, air sac belongs to SK09. Cells positive for influenza A virus antigen stain red in the nucleus and cytoplasm. Brain and pancreas: the centre of the H&E picture contains necrotic debris (loss of architecture, hypereosinophilia, karyorrhexis and pyknosis) close to neuronal cells and exocrine pancreatic cells that express virus antigen (visible in IHC). Air sac: antigen-positive cells largely have their normal morphology, with no associated lesions.

All five skuas that tested negative for H5 HPAIV RNA by RT-qPCR and of which tissues were sampled for histopathology (Hope Bay – SK01 and SK06; Paulet Island – SK29 to 31) were also negative for influenza A virus antigen expression.

#### Co-localization of virus antigen expression and histological lesions

Histological lesions were attributed to H5 HPAIV infection based on their co-localization with virus antigen expression (Fig. 4, fig. S2, table S4). Virus antigen expression in the lungs colocalized with diffuse distension of blood vessels with erythrocytes (SK07 and SK12; fig. S3) and flooding of approximately 10 to 30% of the lumina of all parabronchi and infundibula with eosinophilic homogenous material (edema) and fibrillary eosinophilic material (fibrin) (SK07 only). Virus antigen expression in the brain co-localized with neuronal necrosis and presence of a few heterophils around larger foci of necrosis (Fig. 4). Virus antigen expression in the pancreas co-localized with necrosis of exocrine glandular epithelial cell necrosis with mild heterophil infiltration around larger foci. In both testicle and thyroid gland, it was co-localized with few degenerate and necrotic cells (fig. S2).

#### Microscopic findings associated with avian cholera

All five skuas that tested negative for influenza virus RNA by RT-qPCR, tested positive for *P. multocida* DNA by qPCR (Table 2). Three (SK01, SK06, SK29) of the five skuas had histological lesions in the liver suggestive of *P. multocida* infection (fig. S5, table S4); liver was not sampled in the other two. These lesions were characterized by the presence of randomly distributed, large aggregates of uniformly shaped coccoid bacteria in the liver, in some aggregates co-localized with heterophils and necrotic hepatocytes (SK06 and SK29). In addition, large aggregates of coccoid bacteria, consistent with *P. multocida*, and confirmed by *in situ* hybridization (supplementary text S2), were also present in the blood vessels of other organs sampled in all five birds that tested positive by qPCR (fig. S5).

#### Microscopic findings associated with fungal infection

Histological evidence of fungal infection was present in three skuas (SK01, SK08, SK30) in variable combinations of tissues: trachea, lung, air sac, heart (table S4). Fungal infection was characterized by abundant presence of fungal hyphae (3-6 µm thick, septate, with regular dichotomous branching, consistent with *Aspergillus fumigatus*) (fig. S6). The presence of fungal hyphae was particularly extensive in the lung. Fungal hyphae were co-localized with necrosis and infiltration with heterophils, macrophages and admixed with fibrin and extravasated erythrocytes.

### Virological and bacteriological analyses of fecal samples from live skuas and bacteriological analysis of environmental matrix samples

None of the fecal swabs collected from live skuas at the sites where dead skuas were found tested positive for H5 HPAIV RNA: Beak Island (0/42 fecal swabs), Hope Bay (0/20), Devil Island (0/11), Paulet Island (0/11). For Beak Island, considering that there were approximately 100 live skuas at the time of our visit, it is possible to state that our sample size was sufficient to conclude with 90% confidence that the prevalence of HPAIV in the feces of the live skuas was lower than 5% (assuming test sensitivity of 90% and random sampling) (table S5). None of the 12 fecal environmental swabs collected from skuas at any of the other sites (Elephant Point, D’Urville Monument, Haddon Bay, Yankee Harbour) tested positive for influenza A virus; however, the sample size of feces from skuas at these locations was lower and does not allow for a statement on the prevalence of the virus (table S5).

None of the environmental matrix samples – air (n=4), water (n=10) or soil (n=4) - from Hope Bay, Beak Island or Paulet Island tested positive for *P. multocida* DNA by qPCR, despite the presence of *P. multocida*-positive dead skuas at Hope Bay and Paulet Island.

### Overview of mortality and diagnostic findings in species beyond skuas

During our expedition, we also found and necropsied carcasses of gentoo penguins, Adélie penguins and Antarctic fur seals (*Arctocephalus gazella*) (Table 3). Although we detected low levels of H5 HPAIV RNA in some of the tissue samples from penguins, our follow-up investigations supported other possible causes of death than HPAI. Additionally, we collected brain samples from one snowy sheathbill (*Chionis albus*) and five southern elephant seals (*Mirounga leonina*), but we did not find any evidence of HPAIV infection. These carcasses were in variable states of autolysis and decomposition (also based on GAPDH RT-qPCR data; table S9) ranging from freshly deceased carcasses to skeletal remains with dried-out fragments of brain tissue.

**Table 3.** Overview of diagnoses in gentoo penguins (GP), Adélie penguins (AP), snowy sheathbills (SB), southern elephant seals (ES) and Antarctic fur seals (FS) found dead. All animals tested negative for *P. multocida* DNA by qPCR in all organs.

| Location | Individual | Age class | Decomposition state | Sex | Nutritional condition | HPAI |  | Observations |  | Diagnosis |  |
| --- | --- | --- | --- | --- | --- | --- | --- | --- | --- | --- | --- |
|  |  |  |  |  |  | Brain viral RNA | Viral histo | Macroscopic | Microscopic | Diseases at death in order of severity | Confidence level |
| Elephant Point | GP01-03 | Juv, Ad | Minimal* | nd | P | - | na | Starvation | na | Unknown | Inconclusive |
|  | GP04 | Juv | Advanced | nd | nd | - | na | Unknown | na | Unknown | Inconclusive |
|  | GP06 | Ad | Minimal* | nd | G | - | na | Unknown | na | Unknown | Inconclusive |
|  | ES01-05 | Juv | Advanced | nd | nd | - | na | Unknown | na | Unknown | Inconclusive |
|  | FS01 | Ad | Advanced | nd | nd | - | na | Unknown | na | Unknown | Inconclusive |
| Devil Is. | AP01-02 | Ch, Juv | Moderate/Advanced | M | VP | + | -† | Starvation | nad | 1. Starvation<br>2. HPAI | Medium |
|  | AP03 | Juv | Advanced | nd | VP | -‡ | - | Starvation | nad | 1. Starvation<br>2. HPAI | Medium |
|  | AP04 | Ad | Advanced | F | G | ++ | - | Egg retention | nad | 1. Egg retention<br>2. HPAI | Medium |
|  | AP05 | Ch | Moderate | M | VP | - | na | Starvation | nad | Starvation | Medium |
|  | AP06 | nd | nd | nd | nd | + | na | Unknown | na | Unknown | Inconclusive |
|  | AP07-25 | nd | nd | nd | nd | - | na | Unknown | na | Unknown | Inconclusive |
| D'Urville Monument | GP08-09 | Ch, Juv | Minimal | F | VP | - | na | Starvation | nad | Starvation | High |
|  | GP10 | Ad | Minimal | M | G | ++ | - | Predation, HPAI | Mild histiocytic thyroiditis;<br>nad | 1. Predation<br>2. HPAI | High |
|  | GP11 | Juv | Minimal* | M | VP | - | na | Starvation | nad | Starvation | High |
|  | GP12 | nd | nd | nd | nd | - | na | nd | na | Unknown | Inconclusive |
|  | GP14 | Ch | Minimal | nd | nd | - | na | Unknown | nad | Unknown | Inconclusive |
|  | GP15, AP26 | Juv | Minimal/Moderate | nd | VP | - | na | Starvation | nad | Starvation | High |
|  | SB01 | nd | Advanced | nd | nd | - | na | nd | na | Unknown | Inconclusive |
| Haddon Bay | FS02 | nd | Advanced | nd | nd | - | na | nd | na | Unknown | Inconclusive |
| Paulet Is. | AP60-61, AP63-64, AP67-68 | Ch | Advanced | nd | nd | - | na | nd | na | Unknown | Inconclusive |
|  | AP62 | Juv | Moderate | nd | VP | - | na | Starvation | nad | Starvation | Medium |
|  | AP65 | Juv | Advanced | nd | nd | - | na | nd | na | Unknown | Inconclusive |
|  | AP66 | Juv | Moderate | nd | VP | - | na | Starvation | nad | Starvation | Medium |
| Yankee Harbour | GP16-19, GP21-22 | Ad | Minimal* | nd | nd | - | na | Predation | nad | Predation | Medium |
|  | GP20 | Ad | Advanced* | nd | nd | - | na | Unknown | na | Unknown | Inconclusive |
|  | FS06 | Juv | Minimal* | M | Mo | - | na | Pneumonia¶ | Bacterial pneumonia# | Bacterial infection | High |

Age class: Ch: chick, Juv: juvenile, Ad: adult Sex: M: male, F: female Decomposition state: VP: very poor, P: poor, Mo: moderate, G: good nd: not determined na: not applicable (not sampled) nad: no abnormalities detected \* Scavenged or predated † Tissues were negative for influenza antigen by IHC, and lesions were not detectable. For these cases, ISH was performed on the brain tissue, and few cells were positive. ‡ Animal positive for HPAI viral RNA but not in the brain. ¶ Severe chronic embolic pyogranulomatous pneumonia.

H5 HPAIV RNA was detected by RT-qPCR in 5 of 25 Adélie penguins from Devil Island and 1 of 7 gentoo penguins from D’Urville Monument (table S6). Four of the 5 positive Adélie penguins were confirmed to have a MBCS (PLREKRRKR/GLF). Overall, viral RNA loads in these penguins were lower than in skuas and not detected in all the tissues and swabs. On average, the highest viral RNA load in H5 HPAIV-positive Adélie penguins was detected in the oropharyngeal swabs (10.1 ± 2.1), but comparable results were also found in the brain (9.4 ± 3.2). The positive gentoo penguin showed the highest viral RNA load in the brain (11.8) (table S6). We also detected non-HPAI influenza A viral RNA in tissues and swabs of additional Adélie (n=1, Paulet Island) and gentoo penguins (n=4, D’Urville Monument) (table S7).

Virus antigen expression was not detected in any of the penguin carcasses that were positive for H5 HPAIV RNA. Rare cells in the brain tissues of two Adélie penguins (AP1 and AP2) were positive by *in situ* hybridization (table S8). By histopathological examination, no other clear cause of death was detected for any of these six penguins, which fits with their probable cause of death based on macroscopic examination being either starvation (3/6), egg retention (1/6), or trauma (1/6) (Table 3). For 1/6 penguins (AP6), neither macroscopic assessment was conducted nor histological samples collected.

Nearly all 366 fecal swabs collected from species other than skuas tested negative for influenza A virus RNA (table S5). The exceptions were two fecal swabs collected from apparently healthy individuals: one, which tested positive for H5N1 HPAIV RNA, from a southern giant petrel (*Macronectes giganteus*) at Yankee Harbour, and the other one, which could not be subtyped or pathotyped due to its low viral load, from a gentoo penguin at D’Urville Monument.

## Discussion

In our survey of sites at the South Shetland Islands and Trinity Peninsula in March 2024 (*20*), we diagnosed HPAI as the probable cause of an unusual mortality event involving at least 10 of 46 skuas (including both brown and south polar skuas) found dead at the time of our visit at Beak Island. Evidence supporting this diagnosis included relevant macroscopic observations, high loads of H5 HPAIV RNA, virus antigen expression and typical lesions in the brains and other organs of skuas found dead, combined with lack of evidence of other potential causes of death. These results establish HPAI as a significant cause of mortality of skuas in Antarctica.

Skuas in Antarctica may be both victims and potential vectors of H5 HPAIV, as indicated previously for wild birds in Europe (*27, 28*) and North America (*29*).

Qua victims, the 46 skua deaths at Beak Island attributed to HPAI are undoubtedly far fewer than the actual number of skua deaths from HPAI there during the 2023-2024 breeding season, because of the dead skuas we missed, already had been scavenged by the time of our survey or died outside the site (*30, 31*). In addition, the skua deaths that may have occurred at other unsurveyed sites of the Antarctic Peninsula were missed. For example, high mortality of great skuas attributed to HPAI was seen at colony breeding sites of at least five geographically distant islands in the UK (Hirta, Orkney, Shetland, Flannan, Fair) between June and October of the 2021 breeding season (*32*).

Comparison of the virus distribution, viral loads, and associated pathological changes in the tissues of these south polar skuas and brown skuas and those in the tissue of great skuas with HPAI are informative. In skuas from both hemispheres, neurological clinical signs were associated with HPAIV infections, resulting in (per)acute death of affected individuals (*10, 32*). Despite a limited sample size for postmortem examination, we confirmed systemic HPAIV infections, including a particular tropism for the brain, which suggests that sampling this organ is the most efficient way to detect HPAIV in this species, but also the lungs, based on high viral RNA loads and viral antigen expression. Thus, neurological clinical signs could have occurred before the onset of death, although we did not observe any apparently sick birds alive during our expedition. In this context, body posture appeared to be a limited indicator of HPAI, as in one case (Paulet Island) the skua was more likely to have died of avian cholera. Virus distribution and associated pathological changes in tissues appeared to be more severe and widespread in great skuas (*32*), suggesting south polar and brown skuas may have an even faster fatal outcome from the disease than great skuas.

Local transmission of HPAIV among skuas at Beak Island was likely due to contact with other infected skuas. Our results of virological analysis of skua carcasses support different routes of transmission. Local transmission among skuas may have occurred via intra-species scavenging or cannibalism (based on multiple virus-positive organs that are typically eaten, as well as our observations of skuas scavenging skua carcasses on Beak Island), via contact with contaminated feces (based on viral RNA-positive cloacal swabs) or via contact with contaminated respiratory secretions (based on viral RNA-positive oropharyngeal or tracheal swabs). The freshwater lakes on Beak Island may have been a fomite for HPAIV transmission of skuas bathing together. In a HPAI-associated unusual mortality event of great skuas on Foula (Shetland Isles), the rapid spread of HPAIV among great skuas was considered to have been facilitated by their habit to bathe and socialize at freshwater lochs and pools, where close conspecific interactions occurred (*33*). Alternatively, the higher density of carcasses near freshwater lakes could be related to agonal processes rather than transmission dynamics.

Qua potential vectors, the results of our study do not provide strong evidence either for or against skuas as long-distance vectors of H5 HPAIV. Whereas some skuas seem to die acutely upon infection, we did not follow these infected individuals over time. Therefore, we cannot estimate whether, for example, they can act as vectors of HPAIV in the early stages of infection. Also, we do not know whether other skuas may be infected without showing clinical signs, as occurs in some duck species (*34*). The unusual mortality event of skuas from HPAI in the Weddell Sea area may have started with introduction of H5 HPAIV from elsewhere by an infected skua. This fits with the facts that HPAIV had been detected in skuas on the South Shetland Islands and in other parts of the Antarctic Peninsula in February 2024 (*13*), and we did not observe mortality of other species at Beak Island. Additionally, this is consistent with the more recent detections of H5 HPAIV in skuas on both the east and west coasts of the Antarctic Peninsula, South Georgia, the South Shetland, Falkland, Marion, Crozet and Kerguelen Islands (https://scar.org/library-data/avian-flu). At Hope Bay, breeding brown skuas arrived earlier in the 2023-2024 season than the non-breeding population, which displayed a peak between mid-January and early February, and which was the only population affected by HPAI (Instituto Antártico Argentino, unpublished data). This would also suggest a possible introduction of H5 HPAIV in the Antarctic Peninsula by non-breeding skuas migrating from locations where the virus was already established (e.g. South Georgia and South Shetland Islands). The preferential brown skuas migration for the southwest Atlantic sector of the Southern Ocean and the Patagonian shelf (*16*) also supports this hypothesis. Alternatively, the unusual mortality event may have started with the introduction by an (undetected) infected bird of another species or marine mammal, with subsequent spillover into skuas (*8, 10*).

The results of virological analysis of environmental faecal samples indicated lack of HPAIV circulation in apparently healthy birds at the times of our visits except for a single faecal sample of a southern giant petrel (out of 13 fecal samples collected from this species). The failure to detect HPAIV RNA in environmental fecal samples of apparently healthy birds contrasts with successful detection in samples of birds found dead, particularly in skuas on Beak Island. This is in agreement with previous studies showing that the usefulness of avian fecal samples for HPAI monitoring programs in wild bird populations is limited (*35–37*). Although shedding of infectious virus particles could be assumed for HPAIV-positive individuals found dead, viral loads from cloacal swabs remained low. Furthermore, elaborate and adapted protocols are required for subsequent analyses to allow identification of viral RNA in feces. H5 HPAIV surveillance by environmental fecal sampling, although easy to perform, clearly needs careful consideration in the Antarctic context considering its apparently low reliability to detect HPAIV circulation.

We diagnosed avian cholera rather than HPAI as possible cause of death of skuas at Hope Bay. Evidence supporting this diagnosis includes high loads of *P. multocida* in multiple tissues (indicating systemic bacterial spread), multifocal coccoid bacterial aggregates in the liver colocalized with necrosis and inflammation (indicating severe acute disease from *P. multocida* infection), and no detectable H5 HPAIV RNA in tissues or swabs. Previously, avian cholera had been diagnosed by bacteriological and pathological analyses in brown and south polar skuas found dead in Hope Bay in the breeding seasons of 1999-2000 and 2000-2001 (*23*). These results underline the importance of considering multiple possible causes when investigating unusual mortality events in skuas or other wildlife species, even in the presence of HPAI outbreaks.

Although we found evidence of H5 HPAIV infection in both dead Adélie penguins at Devil Island and a dead gentoo penguin at D’Urville Monument, we diagnosed other probable causes of death, mainly starvation. HPAI was ruled out as cause of death based on the low viral RNA levels by RT-qPCR in tissues and lack of influenza virus antigen expression by IHC and associated lesions by histopathology in the same tissues. There also were two dead skuas at Hope Bay in which low to medium levels of H5 HPAIV were detected in tissues by RT-qPCR, but that possibly or likely died from avian cholera. These results fit with the broad range of disease outcomes in multiple wild bird species infected with HPAIV, from subclinical infection to acute death (*5, 34, 38*). Therefore, the detection of H5 HPAIV RNA by itself in the avian brain does not necessarily mean that HPAIV infection was the cause of death. However, the lack of clinical signs represents a concern for viral persistence in animal populations and potential spread to susceptible species.

We conclude from the detection of H5 HPAIV in five species (brown skua, south polar skua, gentoo penguin, Adélie penguin, southern giant petrel) at four sites at a geographical distance of up to 50 km from each other in our study, together with the detection in other parts of the Antarctic Peninsula by us and others (*8, 13, 20*), that H5 HPAIV already was widespread in the northern part of the Antarctic Peninsula in March 2024, just two months after its first detection in Antarctica (*8*). We also conclude that H5 HPAIV has the potential to cause marked mortality in south polar and brown skuas of the Antarctic Peninsula, up to levels that can potentially cause declines in these small populations. In 1984, it was estimated that there were only 650 breeding pairs of south polar skuas and 150 breeding pairs of brown skuas in the Antarctic Peninsula, out of global populations of 5000 to 8000 breeding pairs of south polar skuas and 7000 breeding pairs of brown skuas (*39*). We are not aware of more recent estimates for the investigated sites, with the exception of Hope Bay. At Hope Bay, the brown skua breeding and non-breeding populations were overall increased between 2014 and 2023, and remained stable between the breeding seasons 2022-2023 and 2024-2025 (39-43 nests and 75-90 individuals, respectively Instituto Antártico Argentino, unpublished data); hence, the mortality at Hope Bay (primarily due to avian cholera) would represent roughly 20% of the local non-breeding population.

Whereas we previously showed that H5 HPAIV RNA was detected in the brain of one of three skuas from Paulet Island (SK30) (*20, 40*), we could not confirm this result hereby. One possible explanation is that since the RNA viral load was low, and given the focal nature of HPAI lesions in the brain (*41*) (as we also showed in the skuas we sampled), one sample might have included a cluster of HPAIV-positive cells, while another one did not. Despite the presence or absence of viral RNA, we diagnosed this animal with a cause of death other than HPAI.

The specific implication of demonstrating that south polar and brown skuas are victims of H5 HPAIV infection is that these two species are at risk of continued high mortality in coming years, especially if the virus is circulating early in the breeding season. This mortality will potentially result in declines of their already small populations, both on the Antarctic Peninsula and elsewhere in their ranges. For comparison, the number of apparently occupied territories of great skuas on Foula (Shetland, UK), the world’s largest breeding colony of this species, was estimated to have declined by about 60-70% due to HPAI (*33*). Similar estimates have also been made for the whole UK (*42*). To be able to assess the effects of HPAI on their populations, it is important to closely monitor as many as possible breeding sites of these species for morbidity and mortality from HPAI. A related priority action is to perform regular population censuses of brown skuas and south polar skuas as frequently as possible in coming years.

The specific implication of evidence for south polar and brown skuas as potential vectors of HPAIV is that they may play a substantial role in further spread of the virus. Breeding colonies of brown skuas are distributed across the Subantarctic region, while those of south polar skuas are distributed along the whole coast of the Antarctic continent. Therefore, they could play a role in spreading HPAIV widely, despite their small numbers. However, relatively little is known about movements of skuas between breeding colonies. To better assess their potential roles as vectors, it is important to learn more about their movement ecology, for example, by attaching miniature data loggers to breeding and non-breeding birds to be able to follow their movements through the environment (*43, 44*).

The general implication of H5 HPAIV being so widespread across the Antarctic Peninsula in several species in the summer of 2023/2024 is that it has gained a foothold in Antarctica and may not only persist but also spread across the continent in coming years. Based on the history of H5 HPAIV in European wildlife in recent years, where the unusual mortality events occurred in different species from one year to the next (EFSA reports 2020 to 2024, from: https://www.efsa.europa.eu/en/topics/topic/avian-influenza), it may be expected that additional species will undergo unusual mortality events from HPAI in Antarctica in the future. Therefore, close avian influenza surveillance by transdisciplinary teams will be needed across Antarctica to be able to record not only virus spread but also diagnose unusual mortality events from HPAI. Such surveillance should include serological screening to understand the role of different species for virus persistence in the Antarctic and be performed according to agreed biosafety guidelines. To be confident that unusual mortality events are caused by HPAI and not by avian cholera or other mortality factors, adequate diagnostic expertise is required. Just as important as recording virus spread and associated mortality, the population sizes of affected species need to be counted in coming years to be able to assess the impact of the virus on wildlife population sizes. This is all the more important for Antarctic wildlife, which is threatened by multiple factors such as global warming, increased scientific activities and infrastructure, increased tourist numbers, invasion of non-native species, overfishing, and pollution (*45*). These avian influenza surveillance and population impact studies fit in a global exercise to assess the impact of this poultry-origin disease on worldwide wildlife populations, and to provide a well-documented basis to take action to prevent future spillover of anthropogenic diseases from livestock to wildlife (*4*).

## Methods Summary

Swabs and tissue samples for molecular analyses, tissue samples for histopathology from animals found dead and environmental fecal and matrix samples were collected from 10 different sites throughout the Trinity Peninsula and Weddell Sea area between March 17th-28th, 2024. Diagnostics analyses were performed to detect influenza A viral RNA (*M1, H5*, HPAI RT-qPCRs and *in situ* hybridization), and influenza A NP viral antigen (IHC). The MBCS was sequenced to confirm the high pathogenicity of the viral detections. In addition, *Pasteurella multocida* diagnostics (*kmt1* qPCR and *in situ* hybridization) was performed to consider avian cholera as an alternative cause of death. The overall diagnosis of each sampled individual was based on a combination of macroscopic and microscopic observations and results of virological and bacteriological diagnostics. A level of confidence (high, medium, inconclusive) was assigned to each diagnosis. A full description of the methods is provided in the supplementary materials.

## Acknowledgments

We thank Ron Fouchier for support in setting up the molecular analyses, Timm Harder and Sanne Thewessen for designing and optimizing the HPAI RT-qPCR protocol, Anne van der Linden for support with the serological analyses, Dirk Höper, Patrick Zitzow, Lukas Wessler and Kristin Trippler for support with the analyses of fecal samples, Jurriaan de Steenwinkel for providing a *P. multocida* positive control, Micaela Carrillo, Diego Torres and Nadia Haidr for field monitoring of the breeding population of skuas at Esperanza/Hope Bay.

