## Supplementary materials for "The expanding avian influenza panzootic: skua die-off in Antarctica"

Materials and Methods

Field work: observations and sample collection

All the samples described in this study have been collected between March 17th and March 28th, 2024, at the following sites:

- Yankee Harbour, Greenwich Island (62°31'55"S, 59°46'32"W)
- Elephant Point, Livingston Island (62°41'21"S, 60°51'58"W)
- Hope Bay, Trinity Peninsula (63°24'16"S, 56°59'49"W)
- Haddon Bay, Joinville Island (63°18'41"S, 55°46'28"W)
- D’Urville Monument, Joinville Island (63°25'36"S, 56°17'13"W)
- Paulet Island (63°34'31"S, 55°46'30"W)
- Beak Island (63°36'49"S, 57°20'50"W)
- Devil Island (63°47'50"S, 57°17'29"W)

The selection of locations we visited was based on sites with a diverse range of wildlife species, especially skua, and previously reported observations of morbidity and mortality. A scouting team was responsible for surveying the territory, recording the number of animals present and observing clinical signs of HPAI in wildlife. The resulting live animal count data do not intend to cover respective population dimensions, but convey the scenario present at the moment of our visit.

Fecal swab samples were collected from apparently healthy individuals by swabbing the dropping/feces and placing it into 1.5 mL NucleoProtect VET buffer (Macherey-Nagel). Autopsies on a subset of dead animals were performed and samples for virological analyses were collected and stored in 1 mL DNA/RNA Shield (Zymo). Samples for histopathological analyses were also collected and stored in 10% neutral buffered formalin.

Water and soil samples were mixed with 0.5 mL Red Blood Cell Lysis Solution (CLB; Promega). Air was collected by filtration through nanofiber filters (*46*) and filters introduced in 2 mL CLB.

All the samples (except tissues for histopathological analyses) were stored refrigerated until the end of the expedition.

Immediate on-site testing for the detection of influenza A virus antigen with the FASTest® AIV Ag test kit (MEGACOR Veterinary Diagnostics, Germany) was conducted for a subset of avian carcasses. Single or multi-organ swabs (table S1) were utilized according to the manufacturer’s instructions; nevertheless, the suboptimal environmental conditions (e.g. low ambient temperature) might have impaired the kit functionality and caused uncertain results in some cases.

All the operations described above were performed with adequate personal protective equipment (PPE).

We applied levels of confidence for our disease diagnoses because we investigated carcasses of free-ranging animals found dead for which we have no disease history knowledge. Additionally, not all carcasses could be investigated with the same tests, either because of lack of time, or because the state of decomposition or scavenging hampered taking a full set of samples. We distinguished three confidence levels: high, medium, inconclusive (table S10). The assigning of these categories was based on i) the samples available for testing, ii) the level of matching of the different test results, and iii) the result of a host housekeeping gene (GAPDH) as internal control for RNA preservation in the carcass. We assigned a category to each necropsied individual for which we made disease diagnosis. GAPDH RNA was detected in all our carcass samples and therefore negative viral and bacterial genes qPCR results were taken into account.

Laboratory analyses: carcass samples

*Nucleic acids extraction*

All samples were stored at –70°C. Tissue samples were homogenized prior virological and bacteriological screenings. RNA was extracted for virological analyses using the High Pure RNA Isolation Kit (Roche) according to the manufacturer’s instructions. For bacteriological analyses, 200 μL of tissue swab or environmental sample supernatant (air, soil, water) and 100 μL of tissue homogenate were mixed with 600 μL of MagNA Pure External Lysis Buffer (Roche) and PBS up to 1 mL. 20 μL of phocine distemper virus (PDV) extraction control was added to each sample (*47*). Total nucleic acids were extracted using MagNA Pure 96 System (Roche).

*M1, H5 and HPAI real time RT-PCR*

The presence of influenza A virus and H5 hemagglutinin (HA) RNA in the samples was determined by duplex RT-qPCR as described previously (*48*).

The presence of a MBCS in the HA sequence was first determined by RT-qPCR. A partial sequence from segment 4 (HA) was amplified using specific primers and probes designed in-house for the detection of HPAI H5 viruses (FW: CCTTGCGACTGGGCTCAG, RV: ATCAACCATTCCCTGCCA, probe1: FAM-AGAAGAAARAGAGGGCTGTTTGGGGCT-BHQ-1, probe2: FAM-AGAAGAAARAGAGGCCTGTTTGGGGCT-BHQ-1). RT-qPCR mix included 20 μL containing 5 μL of RNA, 500 nM of each primer, 250 nM of each probe, 5 μL 4x TaqMan™ Fast Virus 1-Step Master Mix (Applied Biosystems) and nuclease-free water to a final volume of 20 μL. The amplification protocol was 5 min 50°C, 20 s 95°C, 45x (3 s 95°C, 20 s 56°C, 31 s 60°C). Fit point analysis was used to determine the Cq values. Negative and positive controls were included. Samples with Cq>40 were considered negative. The following categories were established based on the M1 viral RNA load (40 – Cq): 0, negative (-); 1 to 10, low (+); 11 to 20, moderate (++); 21 to 40, high (+++).

*MBCS sequencing*

To further confirm the presence of a MBCS, the segment region including the MBCS was amplified by RT-PCR and sequenced. cDNA was first synthesized using 80 nM of non-segment-specific H5N1 primer (AGCRAAAGCAGG), 2 μL dNTPs 10 mM, 20 U RNAse inhibitor and 11.5 μL RNA. This mix was incubated for 5 min at 65°C and then placed on ice for denaturation of RNA secondary structures. 20 U RNAse inhibitor, 1x Superscript IV buffer, 1 μL DTT 0.1 M, 200 U Superscript IV Reverse Transcriptase (Invitrogen) were added to the mix in a final volume of 25 μL. cDNA synthesis was performed with the following parameters: 5 min 25°C, 15 min 50°C, 10 min 80°C. A segment region of 310 bp including the MBCS was amplified using specific primers (J3 and B2a) as described previously (*49*). The reaction mix consisted of 5 μL cDNA, 200 nM of each primer, 5 μL GeneAmp 10X Gold Buffer, 5 μL MgCl_2_ 25 mM, 5 U AmpliTaq Gold DNA Polymerase (Applied Biosystems), 1 μL dNTPs 10 mM and nuclease-free water in a final volume of 50 μL. The RT-PCR amplification was performed with the following parameters: 6 min 95°C, 40x (20 s 95°C, 30 s 50°C, 1 min 72°C), 6 min 72°C. After separation on a 2% agarose gel electrophoresis, bands corresponding to the cleavage site amplicon were purified with MiniElute Gel Extraction Kit (Qiagen) according to the manufacturer’s instructions. The amplicon was then prepared for Sanger sequencing. 2 μL of DNA were amplified using 600 nM of the same primers specific for the cleavage site, 0.5 μL BigDye 5X Sequencing Buffer, 0.5 μL BigDye™ Terminator v3.1 (Applied Biosystems) and water to a final volume of 10 μL. The amplification parameters were the following: 10 s 96°C, 30 s 45°C, 30x (4 min 60°C). The amplification products were then purified on a Sephadex plate PERFORMA V3 (EdgeBio). Sequencing was performed in a 3500xL genetic analyser (Applied Biosystems). Sequences were analysed using BioEdit or SnapGene. A reference sequence will be submitted to GenBank.

*GAPDH and β-actin real time RT-PCR*

To estimate the degree of RNA integrity in the carcass samples, ubiquitously expressed host gene RNAs were amplified by RT-qPCR. From bird samples, GAPDH RNA was amplified using specific primers and probe designed based on Adélie penguin GAPDH gene (Gene ID: 103921100) (FW: CAACCCCCAATGTCTCTGTT, RV: TATATGCCAGGATGCCCTTC, probe: FAM-AAGGCTGCTGCTGATGGGCC). From seal samples, β-actin RNA was amplified using in-house designed primers cross-reactive with seals (FW: GGCATCCATGAAACTACCTT, RV: AGCACTGTGTTGGCATAGAG, probe: FAM-ATCATGAAGTGTGACGTTGACATC). The 20 μL RT-qPCR mix included 5 μL of extracted RNA, 20 μM of each primer, 10 μM of probe, 5 μL 4x TaqMan™ Fast Virus 1-Step Master Mix (Applied Biosystems) and nuclease-free water to a final volume of 20 μL. The amplification protocol included 5 min 50°C, 20 s 95°C, 45x (3 s 95°C, 31 s 60°C). Fit point analysis was used to determine the Cq values. Samples with Cq>40 were considered negative.

*Pasteurella multocida kmt1 real time PCR*

The presence of *P. multocida* in the samples was determined by qPCR. A partial sequence of the *kmt1* gene, which is conserved among all *P. multocida* strains, was targeted using primers as previously described (*50*). The qPCR mix included 5 μL of extracted DNA, 20 μM of each primer, 10 μM of probe, 5 μL 4x TaqMan™ Fast Virus 1-Step Master Mix (Applied Biosystems) and nuclease-free water to a final volume of 20 μL. The amplification protocol was 20 s 95°C, 45x (15 s 95°C, 20 s 53°C, 31 s 60°C).  Fit point analysis was used to determine the Cq values. Negative and positive controls were included. Samples with Cq>40 were considered negative.

To further confirm the presence of the *P. multocida* *kmt1* gene, a subset of samples from different locations were used to amplify and sequence the same amplicon (211 bp) of the qPCR reaction. The PCR mix consisted of 5 μL input DNA, 200 nM of each primer, 1 μL SuperScript III RT/Platinum Taq Mix (Invitrogen), 25 μL 2X Reaction Mix and nuclease-free water to a final volume of 50 μL. The amplification protocol was 6 min 95°C, 40x (20 sec 95°C, 30 sec 53°C, 60 sec 72°C), 6 min 72°C. The PCR product isolation and sequencing steps were the same as described above. The sequences were analyzed using BLASTn against the NCBI GenBank database to identify closely related sequences. The following categories were established based on the *kmt1* DNA load (40 – Cq): 0, negative (-); 1 to 7, low (+); 8 to 14, moderate (++); 15 to 40, high (+++).

*Serology*

Serum samples from dead animals were tested for the detection of antibodies against influenza A viral nucleoprotein (NP) and H5 hemagglutinin (HA) using ID Screen Influenza A Antibody Competition Multi-species and ID Screen Influenza H5 Antibody Competition 3.0 Multi-species (IDvet, Innovative Diagnostics), according to the manufacturer’s instructions.

*Immunohistochemistry (IHC), in situ hybridization (ISH) and histopathology*

IHC for the NP antigen of influenza A virus was performed as described previously (*51*). ISH for the NP segment of influenza A virus was performed as described previously (*52*). ISH for *Pasteurella multocida* was performed as described previously (*52*) using the RNAscope probe B-P.multocida-Pm1-dnaA (Catalog #1267471-C1; BioTechne). The UBC positive control probe worked for the species for which ISH was performed (Adélie penguin, south polar skua, brown skua). 3 µm formalin-fixed paraffin-embedded tissue sections were deparaffinized using xylene, rehydrated using graded ethanol and stained with haematoxylin (Klinipath) and eosin (QPath; H&E) to assess any histopathological changes by light microscopy.

Laboratory analyses: fecal samples

*RNA extraction*

All samples were stored at –70°C. 250 μL of original (first step) or 1:4 diluted (second step) supernatant was treated with Proteinase K for 20min at RT and spun down at 13,000 rpm for 1 min. A pre-clean up by incubation with TRIzol™ LS reagent (Thermo Fisher Scientific, Waltham, MA, USA) and chloroform and RNA extraction was performed as described before (*53*). A heterologous RNA (IC2) was added during the extraction process in 1:10 ratio to the later on elution volume of 110 µL as internal control.

*Real time RT-PCR*

The extracted RNA was used as input for a duplex RT-qPCR screening, targeting IAV M1-gene and the EGFP-control gene of IC2 as described previously (*28*). Positive IAV samples were tested to determine subtype and pathotype. The German National Reference Laboratory for avian influenza RT-qPCR approach has been used for confirmation/exclusion of NP-, H5-, N1 and HPAIV H5-specific targets (*54*). Samples without a clear IC2 signal (Cq>34) have been considered as inhibited and thus not evaluable. Those inhibited samples underwent a second round of previously described clean-up, RNA extraction and RT-qPCR screening, but with a 1:4 diluted original sample material in PBS. If in this second round a sample did not reveal an IC2 signal, it remained not evaluable.

Supplementary Text

S1. Virus antigen expression compared to *in situ* hybridization

Immunohistochemistry for influenza A virus antigen and *in situ* hybridization for influenza A virus RNA were compared in H5 HPAIV-RT-qPCR-positive brain sections of two skuas, one with prominent virus antigen expression (SK07) and another with no virus antigen expression (SK11) in the brain (fig. S4A). Positive *in situ* hybridization was characterized by diffuse pink staining of the entire cell, or coarsely granular pink staining in the nucleus and fine granular staining in the cytoplasm. In the skua with prominent virus antigen expression (SK07), positive *in situ* hybridization co-localized with virus antigen expression; in addition, *in situ* hybridization was positive in approximately 10% more cells in and around the areas expressing virus antigen (fig. S4B). In the skua with no virus antigen expression in the brain (SK11), *in situ* hybridization was positive in large areas of the brain, confirming the RT-qPCR results (fig. S4A).

S2. *Pasteurella multocida* RNA expression in tissues of skuas

Liver tissue of one *kmt1* qPCR-positive skua with *P. multocida*-associated lesions with intralesional bacteria was arbitrarily selected (SK29, +++ by qPCR in lung and brain) and stained by *in situ* hybridization with a probe specific for *Pasteurella multocida*. The ISH showed consistent bright pink staining of the coccoid bacteria present in the H&E serial section of the *P. multocida* ISH, confirming that the bacteria in the liver, which co-localize with lesions, are indeed *P. multocida* (fig. S5B).

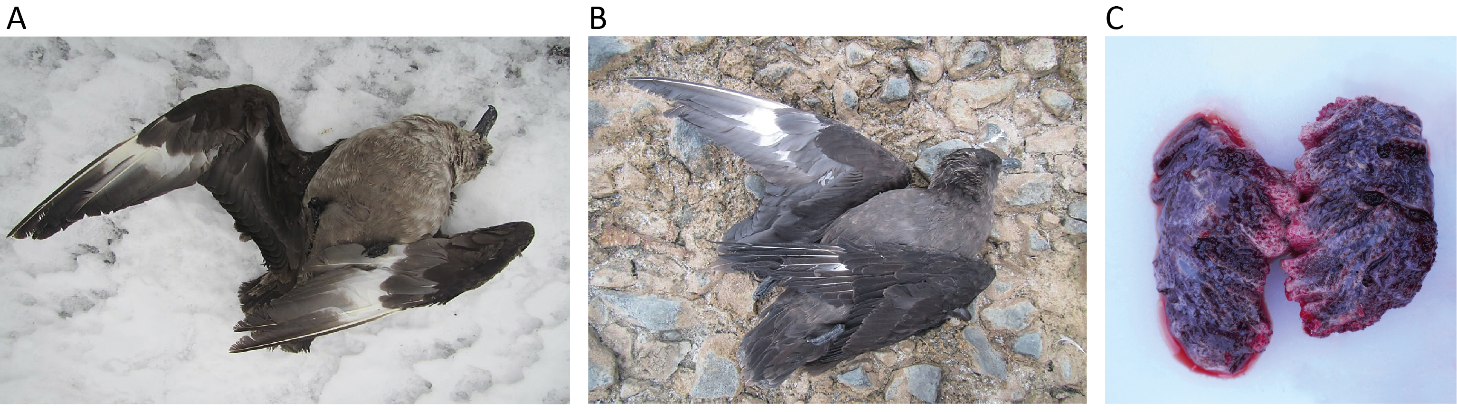

Fig. S1.

**Macroscopic observations in skuas found dead.** (**A**) opisthotonos and wings spread (Beak Island #27), (**B**) torticollis and wings spread (Beak Island #4, SK08), and (**C**) pulmonary oedema (Devil Island #1, SK07).

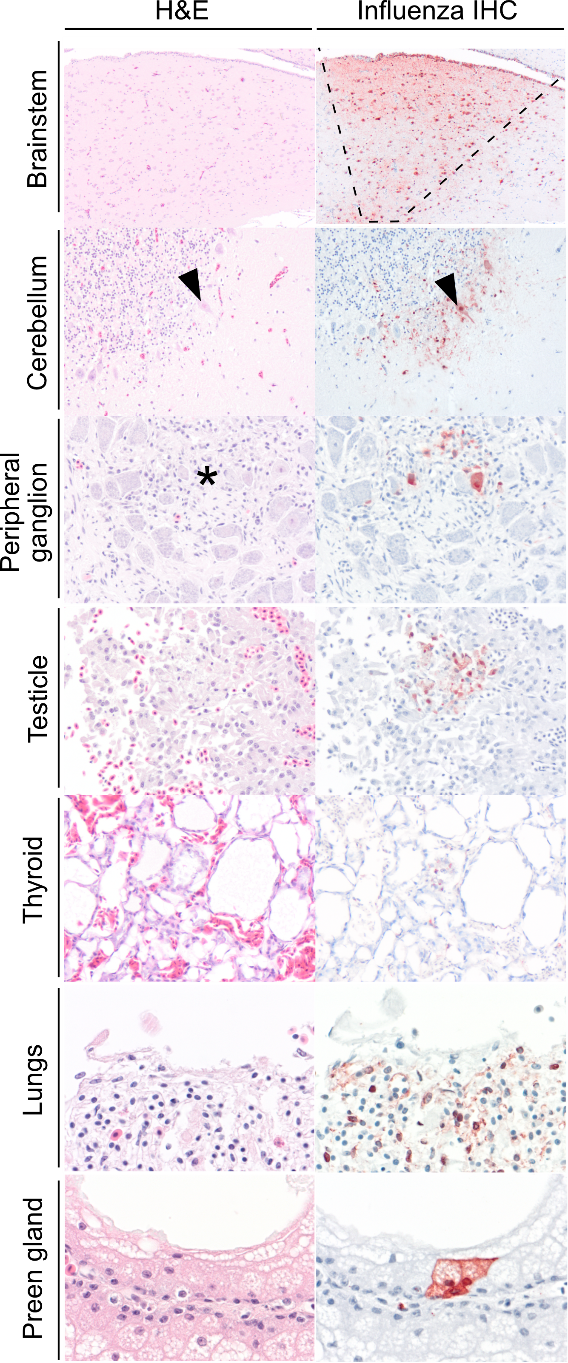

Fig. S2.

**Serial sections of hematoxylin and eosin staining (H&E) and influenza A NP IHC in tissues of skuas found dead.** Preen gland and thyroid were sampled from SK12, all other tissues shown from SK07. Cells positive for influenza virus antigen stain red in the nucleus and cytoplasm. Brainstem IHC shows a large stretch of ependymal cells and a wedge-shaped area of the underlying neuropil expressing virus antigen – indicated by interrupted lines- suggesting entry via the cerebrospinal fluid. Cerebellum shows antigen expression in Purkinje cells, as indicated by arrowhead). The peripheral ganglion shows increased cellularity in the area with neurons expressing virus antigen, area indicated by asterisk. Testicle shows inactive cells lining seminiferous tubules expressing virus antigen, with absence of overt changes to cells and tissue architecture. Thyroid with epithelial cells expressing antigen without overt lesion. Lung shows abundant antigen expression in cells in air capillaries, without apparent lesions in this view. Preen gland shows a small cluster of antigen-expressing epithelial cells lining the lumen of the gland.
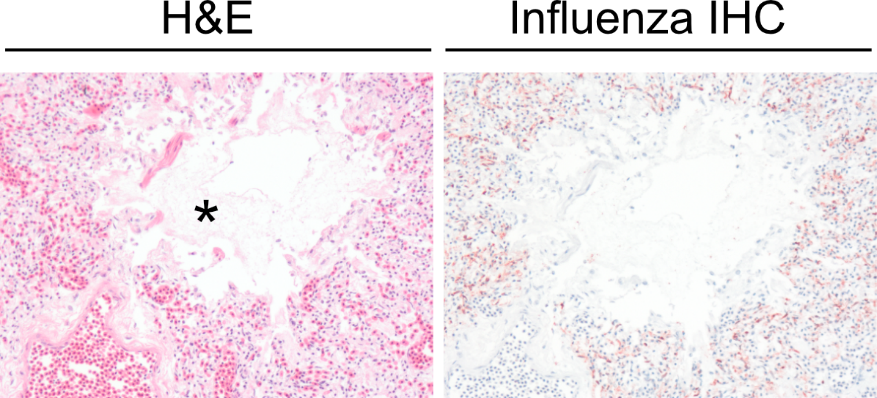

Fig. S3.

**Serial sections of lung of an HPAI-infected skua (SK07).** Approximately 30% of the parabronchus lumen is filled with fibrillar eosinophilic material (fibrin, area with fibrin indicated with asterisk) and homogenous eosinophilic material (oedema). Surrounding capillaries are engorged with erythrocytes (hyperaemia). The serial section shows high levels of virus antigen expression in lung parenchyma.

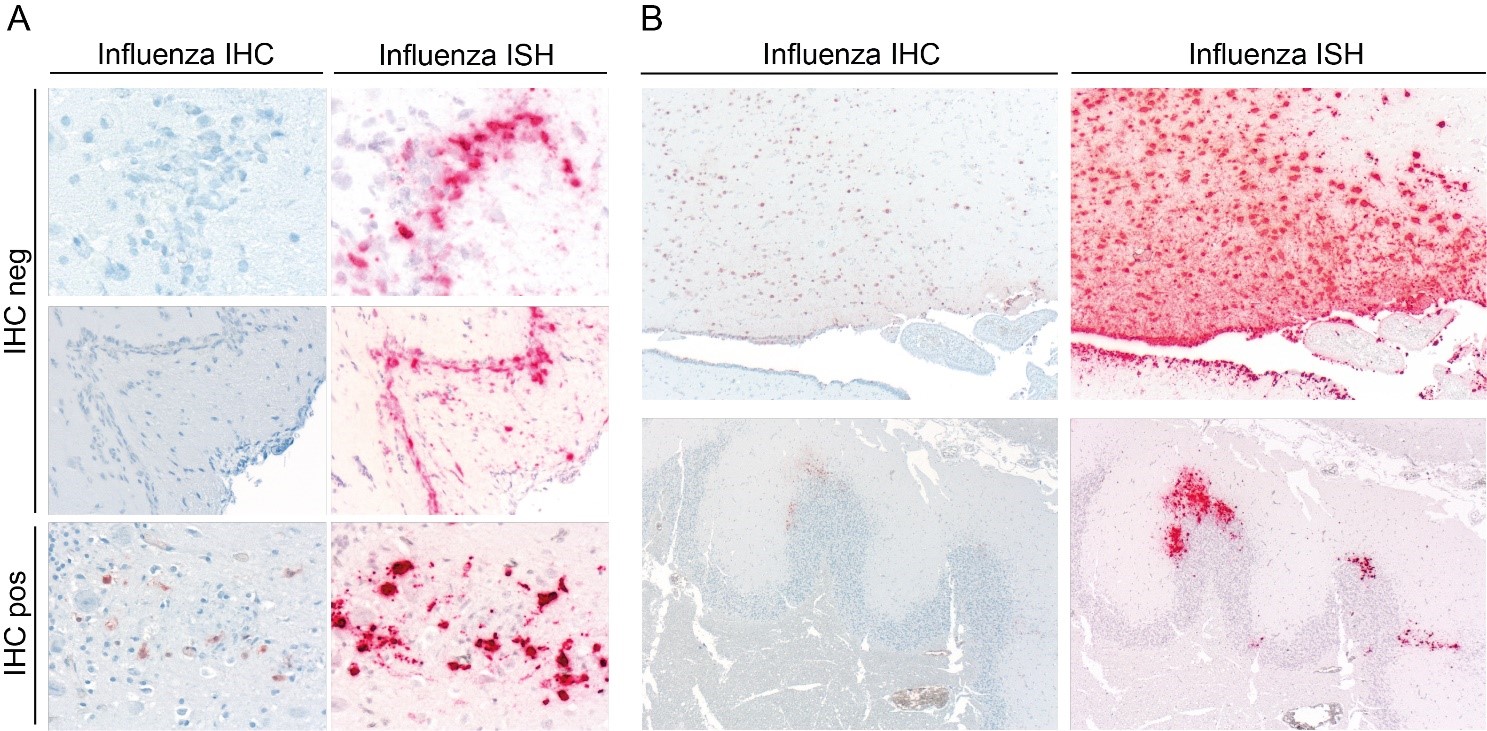

**Fig. S4.**

**Paired influenza A NP immunohistochemistry (IHC) and *in situ* hybridization (ISH) in brain tissues of skuas found dead.** (**A**) Serial sections of skua brain negative (top, SK11) and positive (bottom, SK07) for IHC (viral protein), while both are abundantly positive for ISH (viral RNA), confirming increased sensitivity of ISH for viral detection. (**B**) Top and bottom show two example serial sections at low magnification (40X) of skua brain (SK07)**,** showing positivity overlap between IHC and ISH.

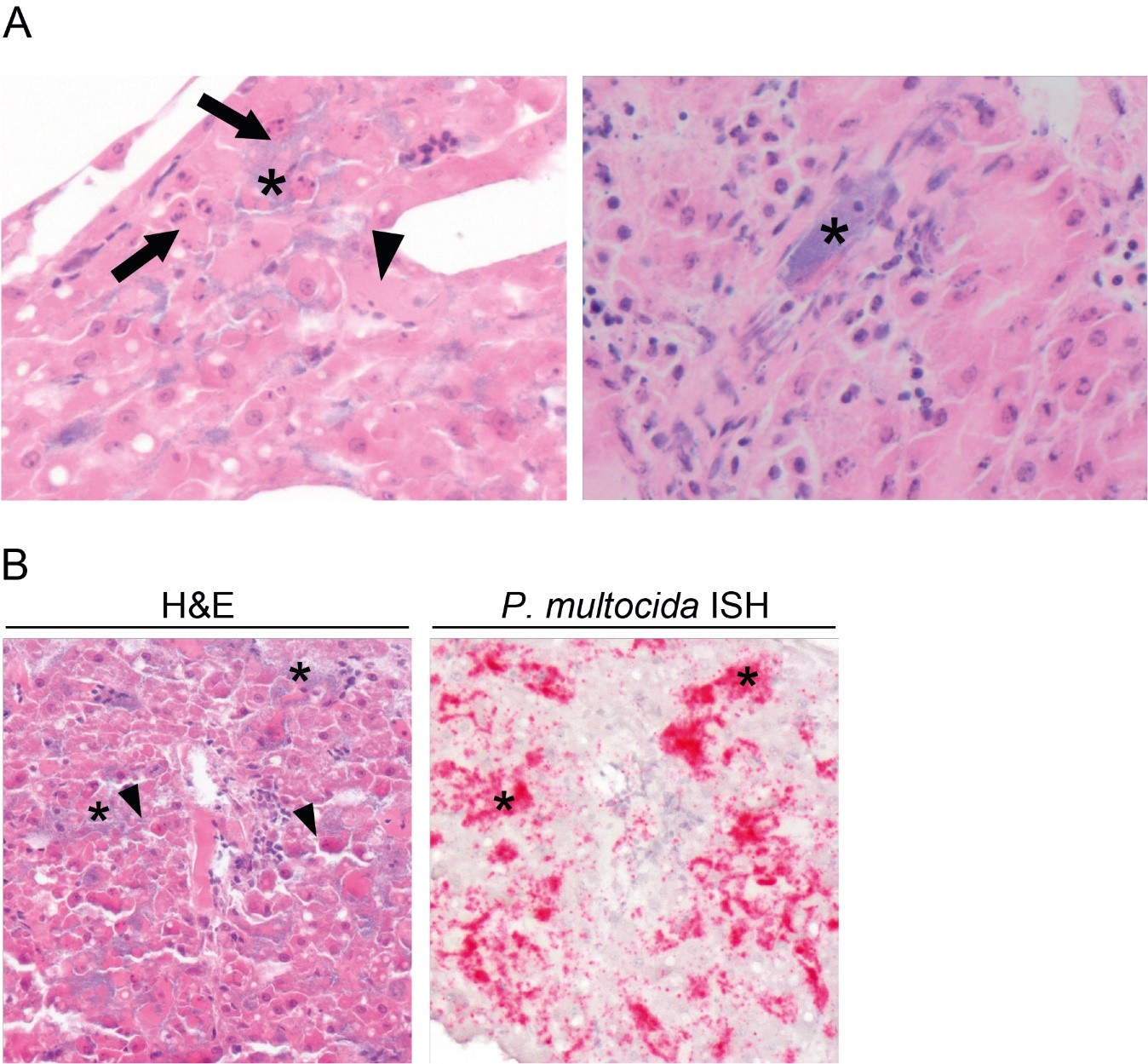

Fig. S5.

Sections of *Pasteurella multocida*-infected liver tissue of skua found dead. (A) H&E panels of bacterial aggregates (asterisk) in liver (left panel) and kidney (right panel) of SK29 (*kmt1*-qPCR-positive). In the liver the bacterial aggregates are associated with heterophils (some indicated by arrows) and degenerating cells (swollen cells with increased eosynophylic cytoplasm, one indicated by arrowhead), and cellular debris, characteristic for necrosis due to *P. multocida* indicated by arrows. In the kidney the bacterial aggregates are present within a blood vessel (asterisk). (B) Serial H&E or *in situ* hybridization (ISH)-stained sections for *P. multocida* of liver tissue of SK29. In the center, the liver tissue architecture is disrupted, and hepatic cords have been replaced with erythrocytes and some cellular debris, characterized by karyorrhexis, swelling and hypereosinophilia of the cytoplasm (arrowheads). Scattered throughout the tissue are abundant aggregates of coccoid bacterial clouds (asterisks), staining positive (bright pink) positive for *P. multocida* by ISH (asterisks).

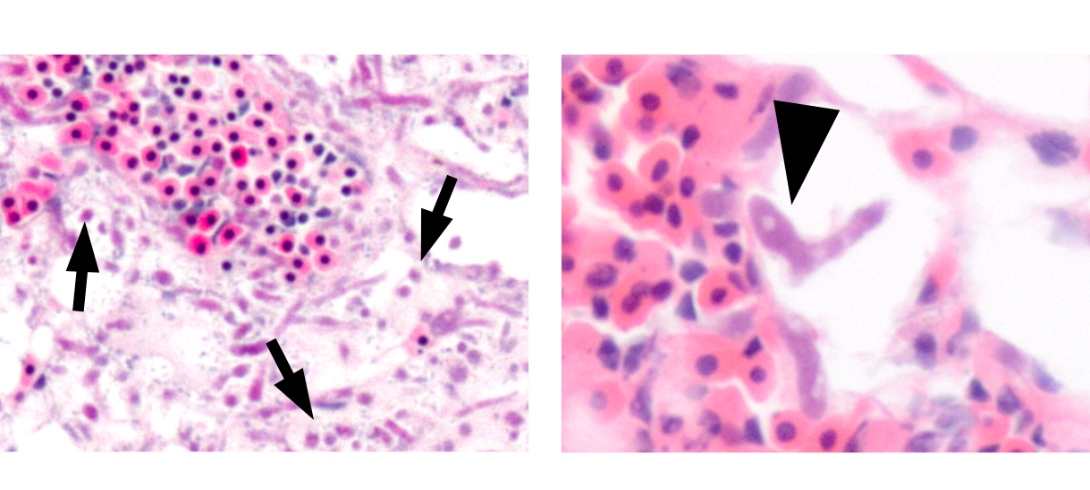

Fig. S6.

H&E panel of fungal hyphae in lung tissue of a skua found dead (SK30). Left panel: fungal hyphae in lung are associated with inflammatory cells (indicated by arrows). Scattered bacteria of different sizes and shapes are consistent with post-mortem overgrowth. Right panel: fungal hyphae are visible as basophilic (purple-blue) structures 3-6 μm thick, septate, with dichotomous branching, consistent with *Aspergillus fumigatus*.
Table S1.

**Macroscopic findings and results of the rapid antigen test screenings of dead skuas**. SK13 to SK17 could not be linked to the carcass ID numbers. The ID numbers from onboard testing, when available, are indicated to compare with previous results (*20, 40*).

| **Location** | **Carcass ID** | **ID** | **Onboard test ID (*40*)** | **Species** | **Age class** | **Nutritional condition** | **State of autolysis** | **Carcass position** | | |  | **Rapid antigen test** | |
| --- | --- | --- | --- | --- | --- | --- | --- | --- | --- | --- | --- | --- | --- |
|  |  |  |  |  |  |  |  | **OP** | **TO** | **WS** |  | **Test result** | **Specimen** |
| **Hope Bay** | 1 | **SK01** | Skua_1_Hope | BRSK | Ad | M | Moderate | na | na | na |  | Negative | O |
|  | 2 | **SK02** | Skua_2_Hope | BRSK | Ad | P | Advanced | na | na | na |  | Negative | O, B |
|  | 3 | **SK03** | Skua_3_Hope | SPSK | Ad | VP | Advanced | na | na | na |  | nd | nd |
|  | 4 | **SK04** | Skua_4_Hope | nd | Ad | VP | Advanced | na | na | na |  | nd | nd |
|  | 5 | **SK05** | Skua_5_Hope | nd | Ad | P | Advanced | na | na | na |  | nd | nd |
|  | 6 | **SK06** | Skua_6_Hope | nd | Ad | M | Moderate | na | na | na |  | Negative | O |
| **Devil Is.** | 1 | **SK07** | Skua_1_Devil | nd | Juv | G | Minimal | yes | no | no |  | Positive | L |
|  |  |  |  |  |  |  |  |  |  |  |  | Uncertain | O, K* |
| **Beak Is.** | 1 | na | na | SPSK | Ad | nd | nd | yes | no | yes |  | nd | nd |
|  | 2 | na | na | SPSK | Ad | nd | nd | no | no | no |  | nd | nd |
|  | 3 | na | na | nd | Ad | nd | nd | yes | no | yes |  | nd | nd |
|  | 4 | **SK08** | Skua_1_Beak | SPSK | Ad | VP | Moderate | no | yes | yes |  | nd | nd |
|  | 5 | na | na | nd | Juv | nd | nd | na | na | na |  | nd | nd |
|  | 6 | na | na | nd | Juv | nd | nd | na | na | na |  | nd | nd |
|  | 7 | na | na | SPSK | Ad | nd | nd | na | na | na |  | nd | nd |
|  | 8 | na | na | nd | Ad | nd | nd | na | na | na |  | nd | nd |
|  | 9 | na | na | nd | Juv | nd | nd | yes | no | yes |  | nd | nd |
|  | 10 | na | na | nd | Juv | nd | nd | yes | no | yes |  | nd | nd |
|  | 11 | na | na | SPSK | Ad | nd | nd | yes | no | yes |  | nd | nd |
|  | 12 | na | na | nd | Juv | nd | nd | yes | no | yes |  | nd | nd |
|  | 13 | na | na | nd | Juv | nd | nd | yes | yes | yes |  | nd | nd |
|  | 14 | na | na | nd | Juv | nd | nd | na | na | na |  | nd | nd |
|  | 15 | na | na | nd | Juv | nd | nd | na | na | na |  | nd | nd |
|  | 16 | na | na | nd | Juv | nd | nd | no | yes | yes |  | nd | nd |
|  | 17 | na | na | nd | Juv | nd | nd | no | yes | yes |  | nd | nd |
|  | 18 | na | na | SPSK | Ad | nd | nd | no | no | yes |  | nd | nd |
|  | 19 | na | na | SPSK | Ad | nd | nd | yes | yes | yes |  | nd | nd |
|  | 20 | **SK09** | Skua_2_Beak | nd | Juv | VP | Mild | no | no | yes |  | Uncertain | L |
|  |  |  |  |  |  |  |  |  |  |  |  | Negative | T, K, I, B |
|  | 21 | na | na | nd | Ad | nd | nd | no | yes | yes |  | nd | nd |
|  | 22 | **SK12** | Skua_5_Beak | nd | Juv | G | Mild | no | no | no |  | nd | nd |
|  | 23 | na | na | nd | Ad | nd | nd | na | na | na |  | nd | nd |
|  | 24 | na | na | nd | Ad | nd | nd | no | no | yes |  | nd | nd |
|  | 25 | na | na | SPSK | Ad | nd | nd | no | no | yes |  | nd | nd |
|  | 26 | na | na | nd | Juv | nd | nd | no | no | yes |  | nd | nd |
|  | 27 | na | na | SPSK | Ad | nd | nd | yes | no | yes |  | nd | nd |
|  | 28 | na | na | nd | Juv | nd | nd | no | yes | yes |  | nd | nd |
|  | 29 | na | na | SPSK | Ad | nd | nd | no | no | no |  | nd | nd |
|  | 30 | na | na | nd | nd | nd | nd | na | na | na |  | nd | nd |
|  | 31 | na | na | nd | Ad | nd | nd | na | na | na |  | nd | nd |
|  | 32 | na | na | SPSK | Ad | nd | nd | no | no | yes |  | nd | nd |
|  | 33 | na | na | SPSK | Ad | nd | nd | no | no | yes |  | nd | nd |
|  | 34 | na | na | SPSK | Ad | nd | nd | no | no | yes |  | nd | nd |
|  | 35 | **SK10** | Skua_3_Beak | nd | Juv | G | Mild | no | no | yes |  | Uncertain | L |
|  |  |  |  |  |  |  |  |  |  |  |  | Negative | K, I, B |
|  | 36 | na | na | nd | Ad | nd | nd | no | no | yes |  | nd | nd |
|  | 37 | na | na | nd | Juv | nd | nd | yes | yes | yes |  | nd | nd |
|  | 38 | na | na | SPSK | Ad | nd | nd | yes | no | yes |  | nd | nd |
|  | 39 | **SK11** | Skua_4_Beak | nd | Juv | G | Mild | no | no | yes |  | nd | nd |
|  | 40 | na | na | SPSK | Ad | nd | nd | yes | no | yes |  | nd | nd |
|  | 41 | na | na | SPSK | Ad | nd | nd | no | no | no |  | nd | nd |
|  | 42 | na | na | SPSK | Ad | nd | nd | yes | no | yes |  | nd | nd |
|  | 43 | na | na | nd | Juv | nd | nd | na | na | na |  | nd | nd |
|  | 44 | na | na | nd | Juv | nd | nd | na | na | na |  | nd | nd |
|  | 45 | na | na | nd | Ad | nd | nd | na | na | na |  | nd | nd |
|  | 46 | na | na | SPSK | Ad | nd | nd | no | no | yes |  | nd | nd |
| **Paulet Is.** | 1 | **SK29** | Skua_1_Paulet | nd | Ad | nd | Moderate | yes | no | yes |  | Negative | L, B* |
|  | 2 | **SK30** | Skua_2_Paulet | nd | Ad | P | Moderate | na | na | na |  | Negative | L, B* |
|  | 3 | **SK31** | Skua_3_Paulet | nd | Ad | nd | Moderate | no | no | yes |  | Uncertain | L, B* |

Species: BRSK: brown skua, SPSK: south polar skua, nd: not determined (species could not be identified with a confidence level of ≥75%).

Age class: Ad: adult, Juv: juvenile

Nutritional condition: G: good, M: moderate, P: poor, VP: very poor

Carcass position: OP: Ophistothonos, TO: torticollis, WS: wings spread, na: not applicable because of scavenging or previous manipulation

Tested specimen: O: oropharynx, T: trachea, B:brain, L: lung, K: kidney, I: intestine

nd: not determined

na: not applicable (not sampled)

*combined swab per specimen and individual

Table S2.

**Results of influenza A virus RT-qPCRs (M1, H5, HPAI) of swabs (s.) and tissues of skuas found dead.** Results are shown as Cq values. The samples from which the MBCS was sequenced are indicated (*). All the sequences had the same MBCS sequence. The ID numbers from onboard testing are indicated in brackets to compare with previous results (*20, 40*).

| **Sample** | **Beak Island** | | | | | | | | | | | | | | | | | | | | | | | | | | | | | | | | | | | | | | |  | **Hope Bay** | | | | | | |  | **Devil Island** | | |
| --- | --- | --- | --- | --- | --- | --- | --- | --- | --- | --- | --- | --- | --- | --- | --- | --- | --- | --- | --- | --- | --- | --- | --- | --- | --- | --- | --- | --- | --- | --- | --- | --- | --- | --- | --- | --- | --- | --- | --- | --- | --- | --- | --- | --- | --- | --- | --- | --- | --- | --- | --- |
|  | **SK08** (Skua_1_Beak) | | |  | **SK09** (Skua_2_Beak) | | |  | **SK10** (Skua_3_Beak) | | |  | **SK11** (Skua_4_Beak) | | |  | **SK12** (Skua_5_Beak) | | |  | **SK13** (Skua_6_Beak) | | |  | **SK14** (Skua_7_Beak) | | |  | **SK15** (Skua_8_Beak) | | |  | **SK16** (Skua_9_Beak) | | |  | **SK17** (Skua_10_Beak) | | |  | **SK02** (Skua_2_Hope) | | |  | **SK03** (Skua_3_Hope) | | |  | **SK07** (Skua_1_Devil) | | |
|  | **M1** | **H5** | **HPAI** |  | **M1** | **H5** | **HPAI** |  | **M1** | **H5** | **HPAI** |  | **M1** | **H5** | **HPAI** |  | **M1** | **H5** | **HPAI** |  | **M1** | **H5** | **HPAI** |  | **M1** | **H5** | **HPAI** |  | **M1** | **H5** | **HPAI** |  | **M1** | **H5** | **HPAI** |  | **M1** | **H5** | **HPAI** |  | **M1** | **H5** | **HPAI** |  | **M1** | **H5** | **HPAI** |  | **M1** | **H5** | **HPAI** |
| **Oropharyngeal s.** | na | na | na |  | na | na | na |  | na | na | na |  | na | na | na |  | na | na | na |  | na | na | na |  | na | na | na |  | na | na | na |  | na | na | na |  | na | na | na |  | 29.4 | 31.1 | 29.9 |  | 34.8 | 33.6 | nd |  | 22.3 | 24.3 | 23.6 |
| **Tracheal s.** | nd | nd | 39.5 |  | 24.1 | 25.1 | 25.2 |  | 23.2 | 25.4 | 25.3 |  | 27.1 | 31.0 | 29.2 |  | 20.2 | 23.3 | 22.0 |  | na | na | na |  | na | na | na |  | na | na | na |  | na | na | na |  | na | na | na |  | na | na | na |  | na | na | na |  | na | na | na |
| **Cloacal swab** | 34.0 | 35.1 | 34.7 |  | 27.8 | 28.3 | 28.3 |  | 26.5 | 27.2 | 27.5 |  | 30.6 | 32.7 | 32.0 |  | 21.5 | 23.8 | 23.2 |  | na | na | na |  | na | na | na |  | na | na | na |  | na | na | na |  | na | na | na |  | 32.6 | 32.3 | 33.8 |  | 39.8 | 36.5 | 37.3 |  | 22 | 25.1 | 23.7 |
| **Brain** | 19.5* | 21.8 | 20.1 |  | 17.8 | 19.6 | 18.6 |  | 18.6* | 20.8 | 20.5 |  | 16.0* | 18.4 | 17.7 |  | 15.0* | 18.0 | 16.9 |  | 12.0* | 14.8 | 14.6 |  | 18.3* | 20.7 | 20.5 |  | 23.7* | 25.4 | 25.2 |  | 17.8* | 21.3 | 19.5 |  | 15.0* | 17.1 | 16.2 |  | 21.4* | 23.2 | 23.0 |  | 34.4* | 37.2 | nd |  | 12.8* | 15.9 | 15.1 |
| **Lung** | 26.6 | 27.6 | 26.7 |  | 21.6* | 23.7 | 23.4 |  | 20.9 | 22.6 | 22.4 |  | 22.0 | 24.2 | 23.9 |  | 13.1* | 16.0 | 15.3 |  | na | na | na |  | na | na | na |  | na | na | na |  | na | na | na |  | na | na | na |  | na | na | na |  | na | na | na |  | 13.3* | 16.1 | 15.1 |
| **Spleen** | na | na | na |  | 25.7 | 26.5 | 25.9 |  | 26.3 | 27.2 | 27.6 |  | 27.8 | 28.6 | 29.6 |  | 13.0 | 16.0 | 15.3 |  | na | na | na |  | na | na | na |  | na | na | na |  | na | na | na |  | na | na | na |  | na | na | na |  | na | na | na |  | 18.2 | 20.4 | 19.3 |
| **Kidney** | 33.4 | 33.1 | 35.3 |  | 30.2 | 30.4 | 31.1 |  | 25.2 | 25.9 | 26.3 |  | 28.4 | 29.0 | 30.0 |  | 14.7 | 18.3 | 17.3 |  | na | na | na |  | na | na | na |  | na | na | na |  | na | na | na |  | na | na | na |  | na | na | na |  | na | na | na |  | 18.7 | 20.6 | 19.5 |
| **Intestine** | 32.1 | 32.3 | 32.3 |  | 28.8 | 29.0 | 29.1 |  | 24.5 | 25.7 | 25.0 |  | 26.1 | 27.9 | 27.5 |  | 16.2 | 18.8 | 18.5 |  | na | na | na |  | na | na | na |  | na | na | na |  | na | na | na |  | na | na | na |  | na | na | na |  | na | na | na |  | 20.3 | 22.0 | 20.9 |
| **Liver** | 31.6 | 32.4 | 32.0 |  | 25.4 | 26.3 | 26.2 |  | 27.3 | 28.1 | 28.6 |  | 29.6 | 31.5 | 31.3 |  | 12.5 | 15.4 | 15.4 |  | na | na | na |  | na | na | na |  | na | na | na |  | na | na | na |  | na | na | na |  | na | na | na |  | na | na | na |  | 19.0 | 21.9 | 20.6 |
| **Feather follicle** | 37.8 | 35.0 | 35.0 |  | 25.0 | 26.0 | 26.0 |  | 30.5 | 30.9 | 31.2 |  | 30.0 | 30.7 | 31.1 |  | 24.4 | 26.4 | 26.3 |  | na | na | na |  | na | na | na |  | na | na | na |  | na | na | na |  | na | na | na |  | na | na | na |  | na | na | na |  | 26.5 | 27.3 | 26.8 |

nd: not detected

na: not applicable (not sampled)

Table S3.

**Results for influenza A virus NP IHC, cell type tropism, and associated histological lesions in tissues of HPAI RNA positive skuas found dead.** The scoring (-, no cells positive; +, few cells positive; ++, few clusters of cells positive; +++, large clusters of cells positive) indicates parenchymal cells of the indicated tissues, unless otherwise mentioned.

| **Sample** | **Devil Island** |  | **Beak Island** | | | | |  | **N° of animals with IHC-positive parenchymal cells/Total n° of animals examined** |
| --- | --- | --- | --- | --- | --- | --- | --- | --- | --- |
|  | **SK07** |  | **SPSK08** | **SK09** | **SK10** | **SK11** | **SK12** |  |  |
| **Brain** | ++ * |  | + | +++* | +++* | - † | +++* |  | 5/6 |
| **Lung** | +++* |  | - | - | - | - | +++* |  | 2/6 |
| **Air sac** | + |  | - (epith. hardly present) | ++ | - | - | + (endoth. only) |  | 2/6 |
| **Pancreas** | ++ (exocrine)* |  | na | - | +* | +* | ++ (exocrine)* |  | 4/5 |
| **Trachea** | - |  | - | - | - | - | ++ (endoth. only) |  | 0/6 |
| **Spleen** | + (endoth. only) |  | na | - | - | - | + (endoth. only) |  | 0/5 |
| **Kidney** | - |  | na | - | - | - | + (endoth. only) |  | 0/5 |
| **Intestine** | + (endoth. only) |  | na | - | - | - | + (endoth. only) |  | 0/5 |
| **Liver** | + |  | - | - | - | - | + (endoth. only) |  | 0/6 |
| **Preen gland** | + |  | na | na | - | - | - |  | 1/4 |
| **Heart** | - |  | - | - | - | - | + (endoth. only) |  | 0/6 |
| **Gonad** | ++* |  | na | na | - | - | + (endoth. only) |  | 1/4 |
| **Thyroid** | + |  | +* | na | na | na | + |  | 3/3 |
| **Adrenal** | + |  | na | na | - | - | + |  | 2/3 |
| **Other** | nad |  | nad | nad | nad | - pituitary and harderian gland | - harderian gland |  | na |

* lesions are co-localized with virus antigen expression

† ISH positive

Endoth.: endothelium, Epith.: epithelium

na: not applicable (not sampled)

nad: no abnormalities detected

Table S4.

**Summary of histological evidence of HPAIV infection, avian cholera (AC), and fungal infection (F) in tissues of skuas found dead.** Emaciation is not clearly visible by microscopy and is not included in this table. All skuas that tested positive for *P. multocida* by qPCR had aggregates of *Pasteurella*-like coccoid bacteria in tissues available for histology, the table only shows the tissues where these bacteria were associated with lesions.

| **Location** | **Individual** | **Brain** | **Trachea** | **Lung** | **Air sac** | **Heart** | **Liver** | **Pancreas** | **Spleen** | **Kidney** | **Intestine** | **Gonad** | **Thyroid** | **Adrenal** | **Preen**  **gland** | **N° of organs evaluated** |
| --- | --- | --- | --- | --- | --- | --- | --- | --- | --- | --- | --- | --- | --- | --- | --- | --- |
| **Hope Bay** | **SK01** | - | F | F | F | F/AC | AC | na | - | - | na | - | na | na | na | 9 |
|  | **SK06** | - | - | - | na | na | AC | na | - | - | - | - | - | - | na | 10 |
| **Devil Is.** | **SK07** | HPAI | - | HPAI | HPAI | - | HPAI | HPAI | HPAI | - | HPAI | HPAI | HPAI | HPAI | HPAI | 14 |
| **Beak Is.** | **SK08** | HPAI | - | F | F | F | - | na | na | na | na | na | HPAI | na | na | 7 |
|  | **SK09** | HPAI | - | - | HPAI | - | - | - | - | - | - | na | na | na | na | 10 |
|  | **SK10** | HPAI | - | - | - | - | - | HPAI | - | - | - | - | na | - | - | 13 |
|  | **SK11** | HPAI | - | - | - | - | - | HPAI | - | - | - | - | na | - | - | 13 |
|  | **SK12** | HPAI | HPAI | HPAI | HPAI | HPAI | HPAI | HPAI | HPAI | HPAI | HPAI | HPAI | HPAI | HPAI | - | 14 |
| **Paulet Is.** | **SK29** | - | na | - | na | - | AC | na | na | - | na | na | na | na | na | 5 |
|  | **SK30** | na | na | F | na | na | na | na | na | na | na | na | na | na | na | 1 |
|  | **SK31** | - | na | - | na | na | na | na | na | - | na | na | na | na | na | 3 |

na: not applicable (not sampled)

Table S5.

**Overview of influenza A virus prevalence in fecal samples collected from apparently healthy wildlife.** The major represented species at the time of the sampling is shown for each location (*). The positive gentoo penguin from D’Urville Monument only tested positive for IAV, while the positive Southern giant petrel from Yankee Harbour was confirmed to be H5N1 HPAIV RNA-positive.

| **Location** | **Species** | **IAV positive/tested** | **Live animals present at the time of visit** | **Minimum sample size** | **Confidence of freedom from HPAI at location**  **(assuming 5% prevalence)** |
| --- | --- | --- | --- | --- | --- |
| **Beak Island** | Skua* | 0/42 | 100 | 37 | 90% |
| **Hope Bay** | Skua* | 0/20 | 60 | 32 | <90% |
| **Devil Island** | Adelie penguin* | 0/28 | 20 | 20 | 99% |
|  | Skua | 0/11 | 15 |  |  |
|  | Weddell seal | 0/1 | 1 |  |  |
| **Elephant Point** | Gentoo penguin* | 0/66 | 500 | 56 | 95% |
|  | Southern elephant seal | 0/2 | 100 |  |  |
|  | Skua | 0/5 | 20 |  |  |
|  | Southern giant petrel | 0/10 | 20 |  |  |
| **D’Urville Monument** | Gentoo penguin* | 1/80 | 300 | 78 | 99% |
|  | Snowy sheathbill | 0/33 | ne |  |  |
|  | Antarctic fur seal | 0/10 | 40 |  |  |
|  | Kelp gull | 0/22 | ne |  |  |
|  | Skua | 0/2 | 10 |  |  |
| **Haddon Bay** | Antarctic fur seal* | 0/20 | 61 | 34 | <90% |
|  | Kelp gull | 0/16 | 2 |  |  |
|  | Skua | 0/1 | 2 |  |  |
| **Paulet Island** | Antarctic fur seal* | 0/15 | 100 | 37 | <90% |
|  | Skua | 0/11 | 30 |  |  |
|  | Snowy sheathbill | 0/1 | ne |  |  |
|  | Weddell seal | 0/1 | ne |  |  |
| **Yankee Harbour** | Gentoo penguin* | 0/60 | 600 | 56 | 95% |
|  | Antarctic fur seal | 0/2 | 35 |  |  |
|  | Skua | 0/4 | 12 |  |  |
|  | Southern giant petrel | **1/3** | 5 |  |  |

ne: not estimated

Table S6.

**H5 HPAIV viral loads (40 – Cq, *M1* gene) in swabs and tissues of Adélie (AP) and gentoo (GP) penguins found dead**. Only animals that tested positive for HPAI RT-qPCR are included.

| **Sample** | **Devil Island** | | | | |  | **D'Urville Monument** |
| --- | --- | --- | --- | --- | --- | --- | --- |
|  | **AP01** | **AP02** | **AP03** | **AP04** | **AP06** |  | **GP10** |
| **Oropharyngeal s.** | 12.2 | 8.1 | 9.8 | na | na |  | 1.8 |
| **Cloacal swab** | 8.6 | 5.9 | nd | na | na |  | 1.8 |
| **Brain** | 10.0 | 9.6 | nd | 12.8 | 5.1 |  | 11.8 |
| **Lung** | 8.1 | 7.8 | 6.6 | 7.9 | nd |  | nd |
| **Spleen** | 8.7 | 5.1 | nd | na | na |  | nd |
| **Kidney** | 5.8 | 4.4 | nd | na | na |  | nd |
| **Intestine** | 5.8 | 6.6 | nd | na | na |  | nd |
| **Liver** | 5.0 | nd | 8.5 | na | na |  | nd |
| **Feather follicle** | 7.3 | 2.4 | nd | na | na |  | nd |

nd: not detected

na: not applicable (not sampled)

Table S7.

**Results of influenza A virus RT-qPCRs (M1, H5, HPAI) of swabs (s.) and tissues of Adélie (AP) and gentoo (GP) penguins found dead.** Results are shown as Cq values. The samples from which the MBCS was sequenced are indicated (*). All the sequences had the same MBCS sequence. Only animals that tested positive for at least one of the three RT-qPCR are included. The ID numbers from onboard testing, when available, are indicated in brackets to compare with previous results (*20, 40*).

| **Sample** | **Devil Island** | | | | | | | | | | | | | | | | | | |  | **D'Urville Monument** | | | | | | | | | | | | | | | | | | |  | **Paulet Island** | | |
| --- | --- | --- | --- | --- | --- | --- | --- | --- | --- | --- | --- | --- | --- | --- | --- | --- | --- | --- | --- | --- | --- | --- | --- | --- | --- | --- | --- | --- | --- | --- | --- | --- | --- | --- | --- | --- | --- | --- | --- | --- | --- | --- | --- |
|  | **AP01** (ADPE_1_Devil) | | |  | **AP02** (ADPE_2_Devil) | | |  | **AP03** (ADPE_3_Devil) | | |  | **AP04** (ADPE_4_Devil) | | |  | **AP06** (ADPE_6_Devil) | | |  | **GP08** | | |  | **GP09** | | |  | **GP10** | | |  | **GP11** | | |  | **GP14** | | |  | **AP68** | | |
|  | **M1** | **H5** | **HPAI** |  | **M1** | **H5** | **HPAI** |  | **M1** | **H5** | **HPAI** |  | **M1** | **H5** | **HPAI** |  | **M1** | **H5** | **HPAI** |  | **M1** | **H5** | **HPAI** |  | **M1** | **H5** | **HPAI** |  | **M1** | **H5** | **HPAI** |  | **M1** | **H5** | **HPAI** |  | **M1** | **H5** | **HPAI** |  | **M1** | **H5** | **HPAI** |
| **Oropharyngeal s.** | 27.8* | 30.0 | 29.6 |  | 31.9* | 34.1 | 35.0 |  | 30.2* | 32.0 | 31.6 |  | na | na | na |  | na | na | na |  | 37 | nd | nd |  | nd | nd | nd |  | 38.2 | nd | nd |  | nd | nd | nd |  | na | na | na |  | na | na | na |
| **Cloacal s.** | 31.4 | 32.7 | 32.9 |  | 34.1 | nd | 39.3 |  | nd | nd | nd |  | na | na | na |  | na | na | na |  | nd | nd | nd |  | nd | nd | nd |  | 38.2 | nd | nd |  | nd | nd | nd |  | na | na | na |  | na | na | na |
| **Brain** | 30.0* | 34.4 | 34.9 |  | 30.4* | 33.3 | 36.6 |  | nd | nd | nd |  | 27.2* | 29.4 | 29.2 |  | 34.9 | 38.7 | 38.8 |  | nd | nd | nd |  | nd | nd | nd |  | 28.2 | 30.9 | 32.8 |  | nd | nd | nd |  | 30.9 | nd | nd |  | nd | nd | nd |
| **Lung** | 31.9 | 34.0 | 33.5 |  | 32.2 | 33.0 | 38.8 |  | 33.4 | nd | nd |  | 32.1 | nd | 38.4 |  | nd | nd | nd |  | 29.4 | 30.8 | nd |  | nd | nd | nd |  | nd | nd | nd |  | 39.8 | nd | nd |  | 37.8 | nd | nd |  | 34.9 | nd | nd |
| **Spleen** | 31.3 | 33.3 | 33.7 |  | 34.9 | 38.0 | 35.9 |  | nd | nd | nd |  | na | na | na |  | na | na | na |  | 36.6 | nd | nd |  | 30.8 | 32.8 | nd |  | nd | nd | nd |  | nd | nd | nd |  | na | na | na |  | na | na | na |
| **Kidney** | 34.2 | 39.9 | nd |  | 35.6 | nd | nd |  | nd | nd | nd |  | na | na | na |  | na | na | na |  | 36.2 | nd | nd |  | nd | nd | nd |  | nd | nd | nd |  | nd | nd | nd |  | na | na | na |  | na | na | na |
| **Intestine** | 34.2 | 39.3 | nd |  | 33.4 | 37.8 | 35.2 |  | nd | nd | nd |  | na | na | na |  | na | na | na |  | nd | nd | nd |  | nd | nd | nd |  | nd | nd | nd |  | nd | nd | nd |  | na | na | na |  | na | na | na |
| **Liver** | 35.0 | 37.1 | 38.7 |  | nd | nd | nd |  | 31.5 | 37.1 | 36.7 |  | na | na | na |  | na | na | na |  | nd | nd | nd |  | nd | nd | nd |  | nd | nd | nd |  | nd | nd | nd |  | na | na | na |  | na | na | na |
| **Feather follicle** | 32.7 | 35.3 | 35.3 |  | 37.6 | nd | nd |  | nd | nd | nd |  | na | na | na |  | na | na | na |  | nd | nd | nd |  | nd | nd | nd |  | nd | nd | nd |  | nd | nd | nd |  | na | na | na |  | na | na | na |

nd: not detected

na: not applicable (not sampled)

Table S8.

**Results for influenza A virus NP IHC, cell type tropism, and associated histological lesions in tissues of HPAI RNA positive Adélie (AP) and gentoo (GP) penguins found dead.** The scoring (-, no cells positive; +, few cells positive; ++, few clusters of cells positive; +++, large clusters of cells positive) indicates parenchymal cells of the indicated tissues, unless otherwise mentioned.

| **Sample** | **Devil Island** | | | |  | **D'urville Monument** | | | |
| --- | --- | --- | --- | --- | --- | --- | --- | --- | --- |
|  | **AP01** | **AP02** | **AP03** | **AP05** |  | **GP08** | **GP09** | **GP10** | **GP14** |
| **Sex** | M | M | F | M |  | F | F | M | nd |
| **State of autolysis** | Moderate | Advanced | Moderate | Advanced |  | Minimal | Minimal | Moderate | Moderate |
| **Nutritional condition** | VP | VP | VP | VP |  | VP | VP | G | nd |
| **Brain** | - * | - * | - | - |  | - | - | - | - |
| **Lung** | - | - | - | - |  | - | - | - | - |
| **Airsac** | - | - | - | na |  | - | - | - | - |
| **Pancreas** | na | na | na | na |  | - | - | - | na |
| **Trachea** | - | - | - | - |  | - | - | - | - |
| **Spleen** | na | na | - | na |  | - | - | - | - |
| **Kidney** | - | - | - | - |  | - | - | - | - |
| **Intestine** | na | na | na | na |  | - | - | - | - |
| **Liver** | - | - | - | - |  | - | - | - | - |
| **Uropythial gland** | na | na | na | na |  | na | na | na | na |
| **Heart** | - | - | - | - |  | - | - | - | - |
| **Gonad** | na | - | na | na |  | - | - | - | na |
| **Thyroid** | - | - | - | - |  | - | - | - | na |
| **Adrenal** | - | - | - | na |  | na | - | - | na |

Sex: M: male, F: female

Nutritional condition: VP, very poor; P, poor; M, moderate; G, good

na: not applicable

nd: not determined

* ISH positive

Table S9.

**GAPDH RNA loads (40 – Cq) in swabs (s.) and tissues of skuas (SK), Adélie (AP) and gentoo (GP) penguins and snowy sheathbills (SB), and β-actin RNA loads (40 – Cq) in swabs and tissues of southern elephant seals (ES) and Antarctic fur seals (FS) found dead.**

| **Location** | **Individual** | **Oropharyngeal s.** | **Tracheal s.** | **Cloacal s.** | **Brain** | **Lung** | **Spleen** | **Kidney** | **Intestine** | **Liver** | **Feather follicle** | **Lymph node** |
| --- | --- | --- | --- | --- | --- | --- | --- | --- | --- | --- | --- | --- |
| **Elephant Point** | **GP01** | 16.8 | na | 14.3 | 21.1 | na | na | na | na | na | na | na |
|  | **GP02** | 16.6 | na | na | 18.5 | na | na | na | na | na | na | na |
|  | **GP03** | 15.9 | na | 12.3 | 20.7 | na | na | na | na | na | na | na |
|  | **GP04** | na | na | 18.0 | 21.5 | na | na | na | na | na | na | na |
|  | **GO06** | 17.8 | na | na | 21.5 | na | na | na | na | na | na | na |
|  | **ES01** | na | na | na | 14.9 | na | na | na | na | na | na | na |
|  | **ES02** | na | na | na | 13.4 | na | na | na | na | na | na | na |
|  | **ES03** | na | na | na | 15.2 | na | na | na | na | na | na | na |
|  | **ES04** | na | na | na | 13.3 | na | na | na | na | na | na | na |
|  | **ES05** | na | na | na | 13.9 | na | na | na | na | na | na | na |
|  | **FS01** | na | na | na | 12.6 | na | na | na | na | na | na | na |
| **Hope Bay** | **SK01** | 17.5 | na | 17.1 | 21.6 | 21.0 | 14.5 | 18.2 | 20.6 | 15.1 | 13.7 | na |
|  | **SK02** | 14.1 | na | 14.3 | 17.7 | na | na | na | na | na | na | na |
|  | **SK03** | 15.4 | na | 15.7 | 16.4 | na | na | na | na | na | na | na |
|  | **SK04** | 17.3 | na | 16.0 | 16.3 | na | na | na | na | na | na | na |
|  | **SK05** | 14.0 | na | 15.6 | 16.3 | na | na | na | na | na | na | na |
|  | **SK06** | 13.3 | na | 19.4 | 19.5 | 17.3 | 13.6 | 17.5 | 20.0 | 17.9 | 14.2 | na |
| **Devil Island** | **SK07** | 19.2 | na | 18.9 | 23.6 | 22.8 | 21.1 | 22.5 | 24.4 | 21.7 | 16.8 | na |
|  | **AP01** | 15.4 | na | 16.3 | 21.1 | 20.3 | 22.6 | 22.9 | 20.6 | 12.3 | 18.5 | na |
|  | **AP02** | 15.4 | na | 18.0 | 21.8 | 19.3 | 8.2 | 19.9 | 18.2 | 11.4 | 20.8 | na |
|  | **AP03** | 15.8 | na | 15.3 | 20.3 | 21.5 | 19.1 | 21.7 | 13.5 | 18.7 | 18.9 | na |
|  | **AP04** | na | na | na | 18.5 | 15.5 | na | na | na | na | na | na |
|  | **AP05** | 14.9 | na | 9.2 | 20.4 | 21.4 | 10.1 | 22.9 | 14.2 | 20.2 | 18.5 | na |
|  | **AP06** | na | na | na | 15.4 | 15.5 | na | na | na | na | na | na |
|  | **AP07** | na | na | na | 16.2 | 15.8 | na | na | na | na | na | na |
|  | **AP08** | na | na | na | 15.1 | 17.3 | na | na | na | na | na | na |
|  | **AP09** | na | na | na | 15.6 | 16.3 | na | na | na | na | na | na |
|  | **AP10** | na | na | na | 15.8 | 17.0 | na | na | na | na | na | na |
|  | **AP11** | na | na | na | 18.7 | 18.8 | na | na | na | na | na | na |
|  | **AP12** | na | na | na | 14.7 | 17.9 | na | na | na | na | na | na |
|  | **AP13** | na | na | na | 15.2 | 18.1 | na | na | na | na | na | na |
|  | **AP14** | na | na | na | 15.7 | 19.4 | na | na | na | na | na | na |
|  | **AP15** | na | na | na | 16.3 | 16.0 | na | na | na | na | na | na |
|  | **AP16** | na | na | na | 15.0 | 17.8 | na | na | na | na | na | na |
|  | **AP17** | na | na | na | 14.4 | 18.1 | na | na | na | na | na | na |
|  | **AP18** | na | na | na | 16.6 | 19.4 | na | na | na | na | na | na |
|  | **AP19** | na | na | na | 16.4 | 18.4 | na | na | na | na | na | na |
|  | **AP20** | na | na | na | 17.3 | 17.1 | na | na | na | na | na | na |
|  | **AP21** | na | na | na | 15.9 | 15.5 | na | na | na | na | na | na |
|  | **AP22** | na | na | na | 16.8 | 17.5 | na | na | na | na | na | na |
|  | **AP23** | na | na | na | 17.0 | 19.0 | na | na | na | na | na | na |
|  | **AP24** | na | na | na | 17.3 | 16.1 | na | na | na | na | na | na |
|  | **AP25** | na | na | na | 17.5 | 18.3 | na | na | na | na | na | na |
| **Beak Island** | **SK08** | na | 14.1 | 17.0 | 20.4 | 20.6 | na | 19.3 | 17.8 | 18.7 | 11.7 | na |
|  | **SK09** | na | 21.5 | 18.0 | 24.1 | 23.7 | 24.9 | 22.2 | 21.6 | 24.6 | 17.5 | na |
|  | **SK10** | na | 18.4 | 17.2 | 23.5 | 23.1 | 20.4 | 23.7 | 23.5 | 23.2 | 19.8 | na |
|  | **SK11** | na | 15.0 | 17.2 | 23.9 | 23.6 | 20.0 | 23.0 | 22.5 | 21.3 | 19.6 | na |
|  | **SK12** | na | 18.7 | 19.8 | 24.0 | 23.3 | 25.0 | 20.5 | 23.6 | 23.4 | 18.9 | na |
|  | **SK13** | na | na | na | 20.9 | na | na | na | na | na | na | na |
|  | **SK14** | na | na | na | 23.3 | na | na | na | na | na | na | na |
|  | **SK15** | na | na | na | 21.8 | na | na | na | na | na | na | na |
|  | **SK16** | na | na | na | 22.3 | na | na | na | na | na | na | na |
|  | **SK17** | na | na | na | 23.6 | na | na | na | na | na | na | na |
| **D'Urville Monument** | **GP08** | 18.9 | na | 9.6 | 23.7 | 23.6 | 25.5 | 23.9 | 24.3 | 24.4 | 21.7 | na |
|  | **GP09** | 21.9 | na | 20.7 | 23.7 | 23.0 | 24.7 | 23.8 | 24.8 | 22.8 | 23.5 | na |
|  | **GP10** | 21.2 | na | 5.9 | 21.1 | 23.4 | 25.6 | 24.0 | 24.0 | 24.0 | 21.4 | na |
|  | **GP11** | 19.7 | na | 20.2 | 23.1 | 21.1 | 24.7 | 24.7 | 24.2 | 24.3 | 24.0 | na |
|  | **GP12** | na | na | na | 24.0 | 22.1 | na | na | na | na | na | na |
|  | **AP26** | na | na | na | 18.4 | 18.3 | na | na | na | na | na | na |
|  | **GP14** | na | na | na | 21.9 | 21.6 | na | na | na | na | na | na |
|  | **GP15** | na | na | na | 22.2 | 22.2 | na | na | na | na | na | na |
|  | **SB01** | na | na | na | 19.6 | na | na | na | na | na | na | na |
| **Haddon Bay** | **FS02** | na | na | na | 14.7 | na | na | na | na | na | na | na |
| **Paulet Island** | **AP60** | na | na | na | 6.9 | 10.7 | na | na | na | na | na | na |
|  | **AP61** | na | na | na | 20.8 | na | na | na | na | na | na | na |
|  | **AP62** | na | na | na | 20.5 | 18.1 | na | na | na | na | na | na |
|  | **AP63** | na | na | na | 20.3 | 21.5 | na | na | na | na | na | na |
|  | **AP64** | na | na | na | 19.8 | 21.7 | na | na | na | na | na | na |
|  | **AP65** | na | na | na | 17.1 | na | na | na | na | na | na | na |
|  | **AP66** | na | na | na | 20.3 | 21.0 | na | na | na | na | na | na |
|  | **AP67** | na | na | na | 19.4 | 21.1 | na | na | na | na | na | na |
|  | **AP68** | na | na | na | 21.2 | 21.4 | na | na | na | na | na | na |
|  | **SK29** | na | na | na | 21.7 | 21.5 | na | na | na | na | na | na |
|  | **SK30** | na | na | na | 20.3 | 19.7 | na | na | na | na | na | na |
|  | **SK31** | na | na | na | 23.6 | 17.5 | na | na | na | na | na | na |
| **Yankee Harbour** | **GP16** | na | na | na | 23.5 | 22.3 | na | na | na | na | na | na |
|  | **GP17** | na | na | na | 21.8 | 21.8 | na | na | na | na | na | na |
|  | **GP18** | na | na | na | 22.8 | 22.1 | na | na | na | na | na | na |
|  | **GP19** | na | na | na | 22.3 | 21.0 | na | na | na | na | na | na |
|  | **GP20** | na | na | na | 20.4 | 21.3 | na | na | na | na | na | na |
|  | **GP21** | na | na | na | 23.0 | 21.0 | na | na | na | na | na | na |
|  | **GP22** | na | na | na | 16.7 | 22.2 | na | na | na | na | na | na |
|  | **FS06** | 19.1 | na | na | 20.8 | 22.3 | na | 23.4 | na | 24.3 | na | 23.2 |

na: not applicable (not sampled)

Table S10.

**Levels of confidence of diagnoses.** Categories were applied to each diagnosis to indicate the confidence level, taking into consideration the lack of disease history for the sampled animals. As GAPDH RNA was detected in all the samples, all the real-time PCR results could be considered regardless of the GAPDH results.

| **Category** | **Samples available for testing** | **Level of matching of results** |
| --- | --- | --- |
| **High** | Sample available both for (RT-)qPCRs and microscopy | (RT-)qPCR result(s) matches  microscopy result |
| **Medium** | Sample available both for (RT-)qPCRs and microscopy | (RT-)qPCR result(s) does not match  microscopy result |
|  | Sample available for (RT-)qPCRs but not for microscopy | Matching of (RT-)qPCR result with microscopy result is not possible |
| **Inconclusive** | (RT-)qPCRs negative or close to the limit of detection, samples for microscopy not available, or no lesions detected that can explain disease leading to death. | As there are no positive results, matching is not applicable |
